# Tau mediated regulation of Rho1- cytoskeletal dynamics in shaping renal tubule development in *Drosophila*

**DOI:** 10.1101/2025.01.22.634261

**Authors:** Neha Tiwari, Madhu G. Tapadia

## Abstract

The present study explored the significance of the microtubule-associated protein Tau in the morphogenesis and physiological processes of Malpighian tubules (MTs) in *Drosophila melanogaster*. Employing genetic manipulation techniques and microscopy, we established that Tau plays a crucial role in MTs development and function. Genetic ablation and RNAi-mediated suppression of Tau led to pronounced structural abnormalities, including cystic formations, non-uniform tubule diameters, and disorganized epithelial architecture. These aberrations are accompanied by perturbations in the cytoskeletal arrangement, compromised cellular polarity, and ionic imbalances. Functional analyses revealed deficiencies in salt homeostasis and fluid secretion within Tau-deficient tubules. Attenuation of Rho1 expression mitigated the structural defects observed in Tau mutants, highlighting the importance of Tau in modulating Rho1-mediated actin-microtubule dynamics suggesting a genetic interplay between Tau and Rho1 GTPase in orchestrating tubule morphogenesis The phenotypic parallels between Tau-deficient MTs and mammalian models of polycystic kidney disease implies an evolutionarily conserved function of Tau in tubular organ biology.

**Graphical Abstract:** 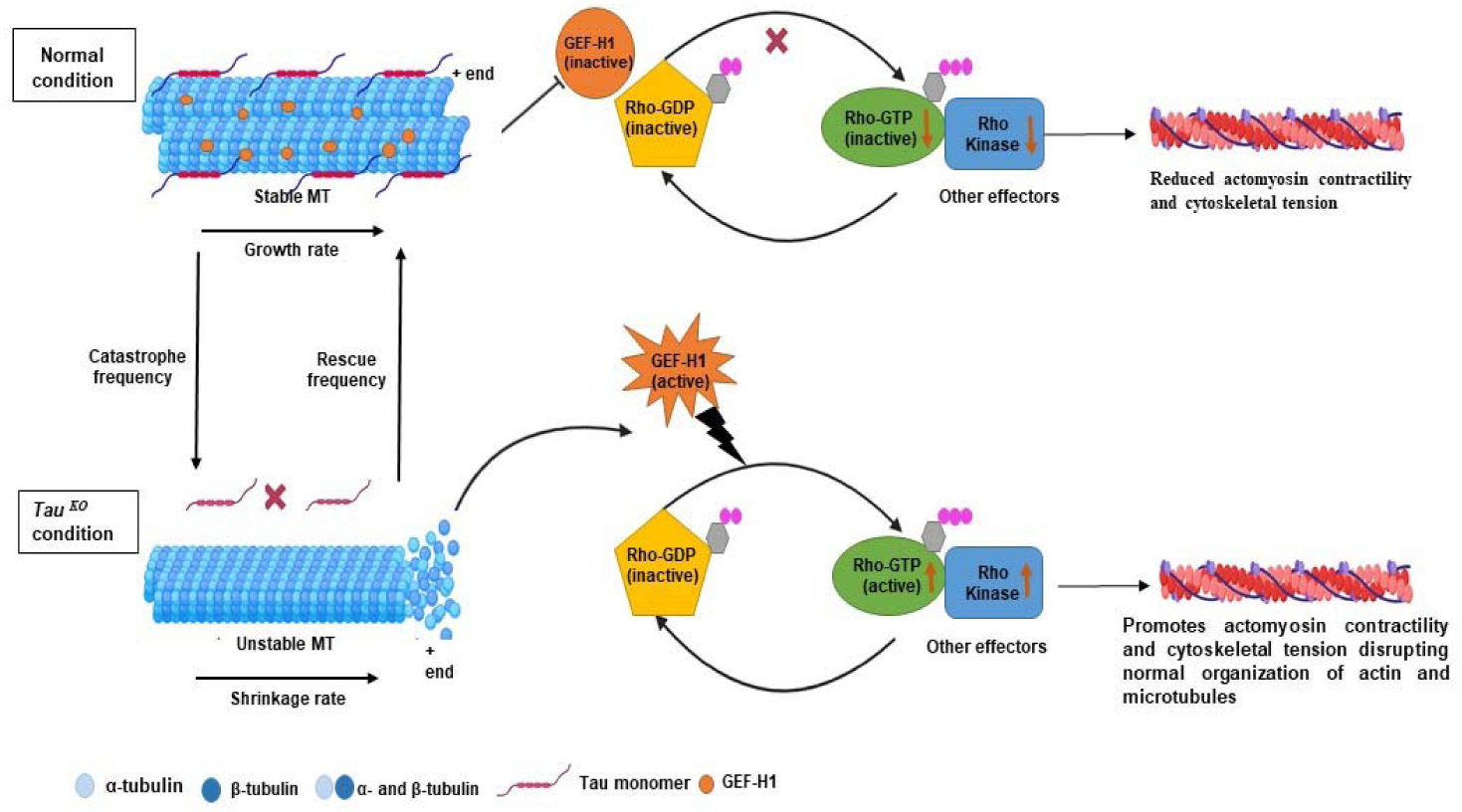

## 1. INTRODUCTION

Tubulogenesis, the development of tubular structures, involves intricate interactions between diverse cell types and environmental signals. This process involves multiple regulatory elements, including transcription and growth factors and their receptors, the modulation of membrane and cytoskeletal dynamics, and protein trafficking (Dressler, 2002; Uv et al. 2003). Appropriate dimensions and morphology of epithelial tubes are critical for the optimal function of organs such as the kidney. Any deviation from normal development can result in functional impairment or other pathological conditions (Dell et al. 2004).

The Malpighian tubules (MTs) are specialized excretory organs in *D. melanogaster* that play a crucial role in maintaining osmoregulation and waste elimination. Functionally analogous to the kidneys in vertebrates, such as humans (Millet-Boureima et al., 2018; Dow and Romero, 2010), they are an ideal model for studying developmental events, such as cell shape changes, adhesion, proliferation, differentiation (Hatton-Ellis et al., 2007; Wan et al., 2000), cellular behavior, and genetic and environmental factors. Disruption of these events causes malformations or lethality (Jung et al., 2005; Denholm, 2013). MT development begins during embryogenesis and is fully functional by the first-instar larval stage (Beaven and Denholm, 2010, Beaven and Denholm, 2018). At the 14-15 stage of development, tubule elongation occurs through cell rearrangement (Abrams et al. 2003; Brooke et al. 2003), and convergent extension movements (Skaer, 1996; Dow, 2012; Beaven and Denholm, 2018; Bunt et al., 2010).

MTs are divided into four regions, ureter, main, initial, and transitional, each with specific excretory and osmoregulatory functions. The initial segment secretes ions and metabolic waste, the main segment (the longest) transports fluid and ions, and the transitional segment links the main segment to the ureter, which delivers the processed fluid to the hindgut for excretion. The epithelial lining of tubules is primarily composed of two cell types: principal cells (PCs) and stellate cells (SCs). PCs play crucial roles in ion transport and contain a high-density population of ion channels that control potassium and sodium levels, which are critical for maintaining osmotic balance and fluid homeostasis (Terhzaz et al., 2006; Tapadia et al., 2011; Evans et al., 2005). Any imbalance in potassium levels is countered by the chloride movement of SCs via chloride channels. These channels drive water transport from the hemolymph through the aquaporin DRIP channels and paracellular pathways (O’Donnell et al., 1983; Kaufmann et al., 2005). Additionally, renal and nephritic stem cells (RNSCs), or “tiny cells,” are also seen in the ureters and lower tubules. These cells migrate from the midgut during metamorphosis, and contribute to tissue repair (Takashima et al. 2013; Li et al. 2014, 2015; Singh et al. 2007).

Recent studies have shown the function of the microtubule-associated protein Tau in cellular phenomena, particularly its role in stabilizing microtubules in neurons, a function critical in neurodegenerative diseases to maintain proper cytoskeletal dynamics. Hyperphosphorylation of Tau may lead to the formation of neurotoxic tangles, which are hallmarks of Alzheimer’s disease and other tauopathies (Avila et al., 2004; Gotz et al., 2019; Goedert et al., 2019). Evidence suggests that Tau protein affects the behavior of non-neuronal tissues, including renal function (Liu et al., 2021; Kim et al., 2019). Although its role in the nervous system is well documented, its involvement in organ development and functionality in non-neuronal tissues, such as MTs, remains under-explored.

Our experiments provide compelling evidence that Tau plays an important role in MTs development and function. Preliminary analysis of Tau knockout (*tau ^KO^*) mutants showed severely disorganized MTs, characterized by aberrant tubular budding and branching, suggesting that Tau is essential for MTs structural integrity. Morphological disruption is characterized by an increase in both the number and length of PCs and SCs. Rho1GTPase and actin cytoskeleton, which are critical for epithelial cell function, were also disrupted due to the absence of Tau protein, which leads to irregularity in cellular polarity, protein expression, localization, and cytoskeleton organization. This disruption of cellular architecture is further exacerbated by abnormal ion channel expression, leading to a significant reduction in luminal diameter, which likely contributes to impaired fluid transport. Furthermore, functional assays revealed diminished MTs functionality in *tau ^KO^* mutants, as these flies exhibited an impaired ability to regulate ion balance and osmotic stress compared to wild-type flies.

By analyzing Tau expression, localization, and interactions within MTs, this study aims to uncover its significance beyond the nervous system in the MTs of *D. melanogaster*. This study suggests a potential role for Tau in human kidney disorders, enhancing our understanding of its broader effects and paving the way for future research into its role in kidney development and disease.

## 2. RESULTS

### 2.1 Tau Expression and Localization in Malpighian Tubules

To characterize the function of Tau in MTs, we used *tau ^KO^/tau ^KO^* (Fig. 1B, E), a line with a selective deletion of the E2 to E6 region (Fig. 1H) of the Tau gene (exon numbering based on FlyBase data for tau-RA transcript), thereby avoiding mutations in surrounding genes, such as rps10a and mir-1001. To first investigate the reduction of Tau in *tau ^KO^/tau ^KO^* condition we performed immunohistochemistry on 3^rd^ instar larval MTs using anti-tau antibody. Tau expression was completely absent in the cytoplasm of both PCs and SCs, confirming the loss of Tau in these genetic backgrounds.

**Figure 1.**
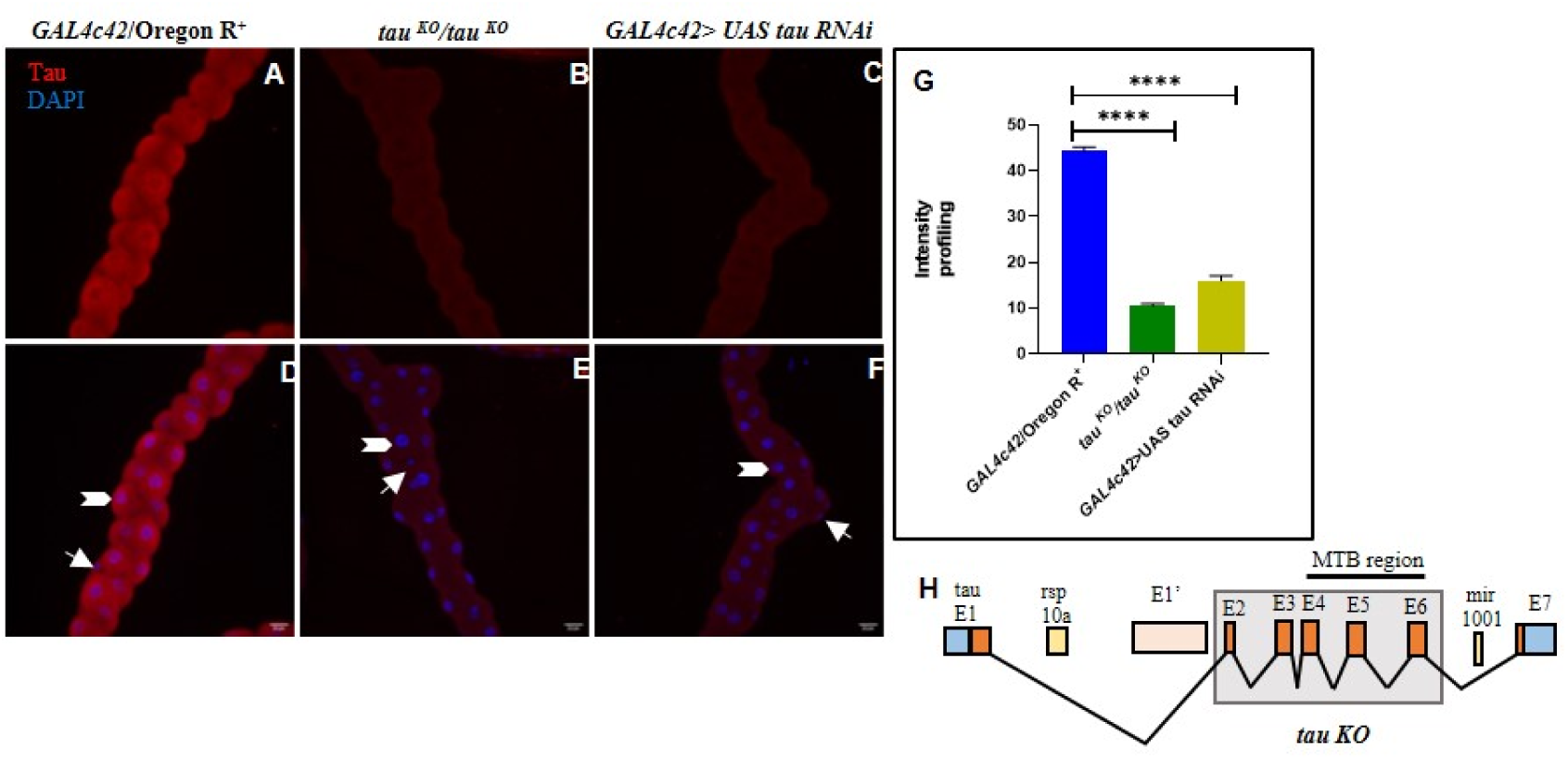
Tau localization in Malpighian tubules under control and Tau-depleted conditions. (**A, D)** In *GAL4c42*/Oregon R^+^ control tubules, Tau is strongly localized within the cytoplasm of principal cells (PCs, shown with arrowhead) and stellate cells (SCs, shown with arrow). **(B, E)** In *tau^KO^ /tau^KO^* tubules, Tau expression is completely absent in both PCs and SCs (shown with arrowhead and arrow), indicating successful knockout of the Tau protein. **(C, F)** Similarly, in *GAL4c42>UAS tau RNAi* tubules, Tau localization is abolished, confirming efficient RNAi-mediated knockdown. Images **A–C** represent Tau immunostaining (red), while **D–F** shows DAPI staining (blue) of the same tubule. Scale bars: 20 µm. **(G)** is a bar diagram showing mean (SE±) intensity profiling of *Tau* of Malpighian tubules in *tau^KO^* and *RNAi* compared to *GAL4c42*/Oregon R Statistical analysis was done using t test, asterisk indicates P ≤ 0.0001. **(H)** Simplified representation of the Drosophila tau gene based on Flybase data for tau-RA/RB transcripts, and the surrounding annotated genes. E1’ is indicated as an alternative exon that is part of the tau-RG transcript. Exons 2 to 6 of the tau gene, including the microtubule binding (MTB) region, were removed by homologous recombination.

We also reduced the expression of Tau using *UAS tau RNAi* driven in MTs cell specific manner by *GAL4c42,* employing Brand and Perrimon method (1993), and compared the pattern of expression in wild type (Oregon R^+^) *Drososphila.* In Oregon R^+^ control tubules, Tau was strongly localized within the cytoplasm of PCs and SCs (Fig 1A, D), whereas in *GAL4c42 > UAS tau RNAi* expression was strongly reduced (Fig. 1C, F), To validate these observations, reverse transcriptase-polymerase chain reaction (RT-PCR) was performed on MTs extracts from *GAL4c42*/Oregon R^+^ (control), *tau ^KO^/tau ^KO^*, and *GAL4c42> UAS tau RNAi* knockdown 3^rd^ instar larvae (Fig. S1A). In the *tau ^KO^/tau ^KO^* condition, Tau mRNA expression levels were significantly reduced by 96% compared to the control, whereas a small tau-RE transcript may still be expressed in the *tau ^KO^/tau ^KO^* line, which likely encodes a 10 kDa Tau-PE polypeptide. However, since this polypeptide lacks the microtubule-binding region (MTBR), it is unlikely to interact with microtubules (Burnouf et al., 2016), while the *GAL4c42>UAS*-*tau RNAi* group also exhibited a significant reduction in Tau mRNA levels (70%) compared to the control, confirming effective knockdown. These data confirm the immunohistochemistry results, showing that Tau is both expressed and localized in the tubules and that its levels can be modulated through genetic manipulations. Together, these findings demonstrate that Tau is expressed in MTs and is particularly concentrated in the cytoplasm and apical regions of PCs and SCs, likely playing a role in maintaining cellular structure and function.

### 2.2 Malpighian Tubule Morphological Deformities Due to Tau Dysfunction

This initial observation confirmed Tau’s presence in the MTs, so we next sought to investigate its role in MTs development. To investigate its functional significance, we selected Tau knockout (*tau ^KO^/tau ^KO^*) mutants and utilized Tau RNAi for further experiments. In Tau-deficient models (both knockout and RNAi-induced knockdown), MTs exhibited clear morphological deformities compared to the controls. Under normal conditions MTs presented a characteristic elongated, coiled structure with uniform diameters throughout the length of larval and adult stages with well-organized cell arrangement (Figure 2A, D). However, *tau ^KO^/tau ^KO^* (Figure 2 E, F) and *GAL4c42*>*UAS tau RNAi* (Figure 2 B, C) tubules displayed multiple cysts in both tubules and an irregular, disorganized arrangement of epithelial cells, indicating a failure in proper tissue architecture. There was a visible loss of uniform diameter in almost all the mutant tubules. Since MTs consist of anterior and posterior tubules, we analyzed whether there was a bias in tubular defects. Statistical analysis revealed that the 30% anterior pair of tubules was defective, 16% posterior pair of tubules was defective, and in 54% larvae, both pairs of tubules were defective, which was higher in comparison to only one pair of defective tubules, which was the anterior pair (Figure 2G).

**Figure 2.**
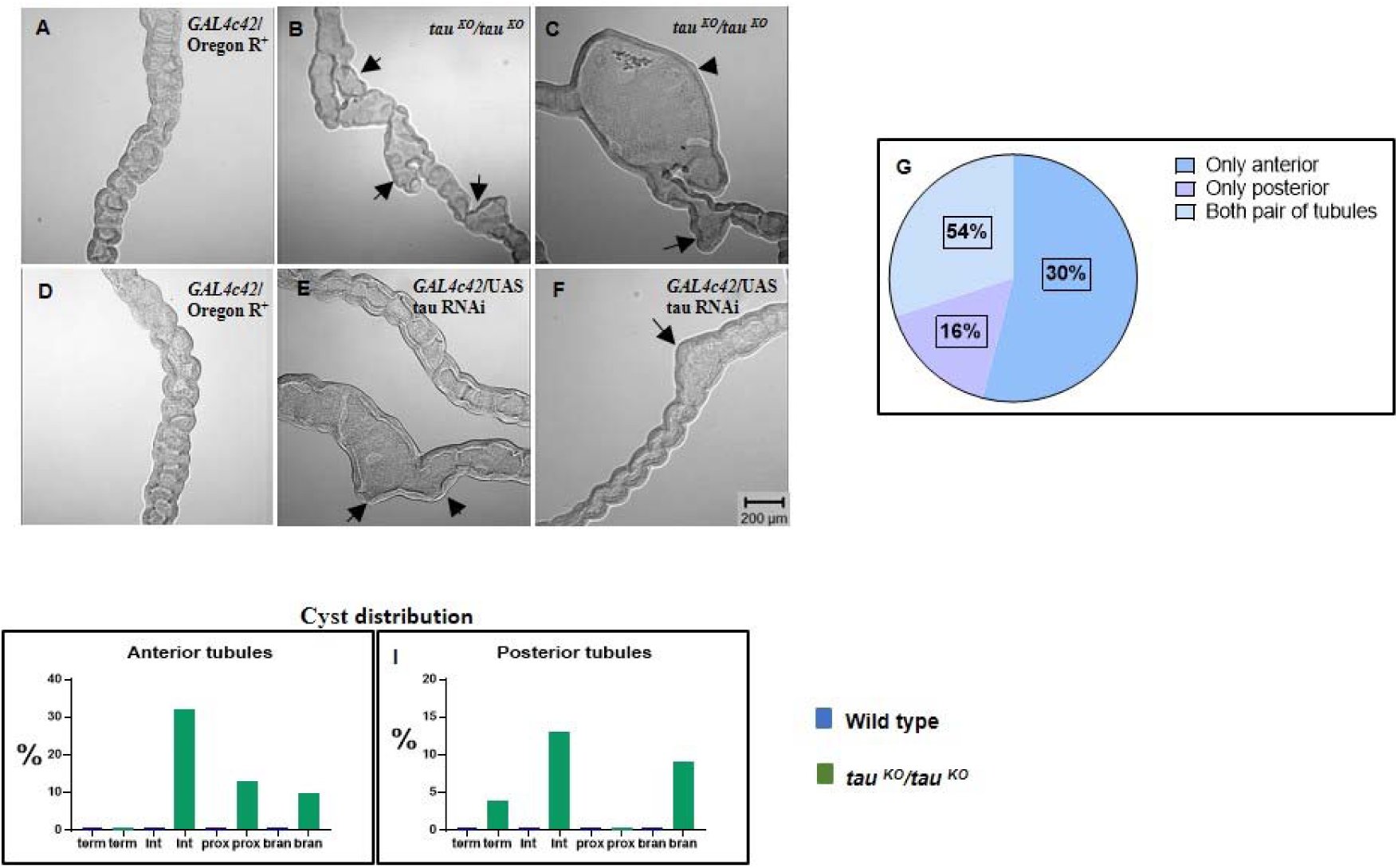
Tau is necessary for proper Malpighian tubule formation. Differential interference contrast images of third instar wild-type MTs **(A)** showing elongated tubules with uniform diameter. *tau ^KO^/tau ^KO^* **(B, C)** and *GAL4c42*/UAS tau RNAi **(E, F)** mutants displayed visible loss of uniform diameter and presence of cysts like structures and twists (arrows) scale bar, 200 μm. **(G)** Proportion of all the four tubules affected in comparison to either the anterior or posteriors tubules in *tau ^KO^/tau ^KO^* (n = 50). **(H, I)** Shown are the percentages of tubules affected in the terminal (term), intermediate (int), and proximal (prox) regions, as well as the observed extra tubular branching (bran). Anterior and posterior tubules were scored separately.

Additionally, extra-tubular budding and branching were observed exclusively in the *tau ^KO^/tau ^KO^* mutant tubules (Figure 2 B and C). To better characterize the defects and progression of the phenotype over time, we scored the incidence of cystic deformations in MTs dissected from 50 3^rd^ instar larvae. Both pairs of MTs displayed cysts, especially in the terminal and intermediate regions (Figure 2 H, I), which was reminiscent of polycystic kidney disease (PKD), which preferentially affects the terminal section of the nephron (Baert, 1978; Grantham JJ et al., 1987) Similar to PKD, the *tau ^KO^/tau ^KO^* mutant tubules also showed extra branching. Tubular cyst number in control and *tau ^KO^/tau ^KO^* conditions categorized into 0, 1, 2, and 3+ groups is shown (Figure S 2A). No tubules were observed in the anterior tubules, with 0 cysts. In total, 44% of the tubules had one cyst, 34% had two cysts, and 22% had three or more cysts. In contrast, in the posterior tubules, 6% of the tubules had 0 cysts, 56% had 1 cyst, 24% had 2 cysts, and 14% had 3 or more cysts.

### 2.3 Ultrastructural and Elemental Insights into Malpighian Tubule

Scanning Electron Microscopy (SEM) revealed distinct structural differences between *GAL4c42*>Oregon R^+^, *tau ^KO^/tau ^KO^*, and *GAL4c42>UAS tau RNAi* MTs. *GAL4c42*>Oregon R^+^ tubules displayed smooth epithelial surfaces with well-defined cellular boundaries, indicating organized tubule architecture (Figure 3 A and A’). In contrast, *tau ^KO^/tau ^KO^* and *GAL4c42>UAS tau RNAi* tubules exhibited a rough, irregular surface with disrupted cellular integrity, suggesting structural deformities (Figure 3B-C’). These morphological alterations align with the observed phenotypes of cytoskeletal disorganization and impaired tubule function in Tau mutants.

**Figure 3:**
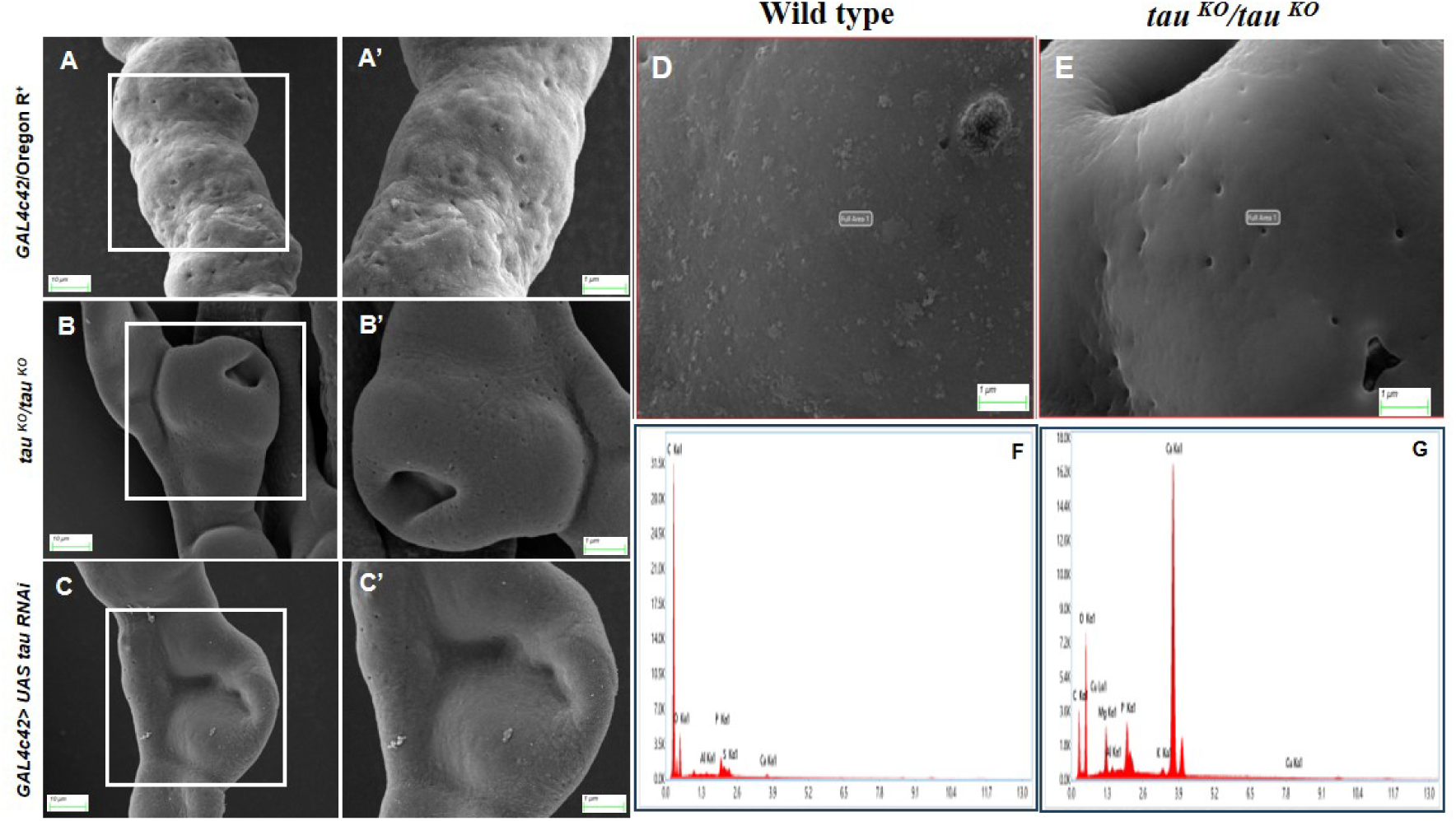
SEM images of Malpighian tubules showing surface morphology differences. **(A)**SEM images of *GAL4c42*>Oregon R^+^ MTs showing smooth surface morphology, clear tubular structure, and uniform cell arrangement, while in *tau ^KO^/tau ^KO^* condition **(B)** MTs displayed disrupted surface morphology, irregularities in cell arrangement, and deformations along the tubule length and in *GAL4c42>tau RNAi* **(C)** MTs exhibited phenotypes similar to *tau ^KO^*, including surface irregularities and structural deformities. **(D)** EDAX spectra of wild-type MTs showing the normal elemental composition, including peaks **(F)** for key elements such as carbon, oxygen, and trace metals, while in *tau ^KO^/tau ^KO^* condition **(E)** EDAX spectra revealed deviations in elemental composition compared to wild-type, with altered peak intensities **(G)** suggesting potential deposition of abnormal aggregates or changes in tissue composition. A’, B’, and C’ are magnified images of the squared boxes in A, B, and C. Scale bar in A, B, C-10 μm and in A’, B’, C’, D and E-1 μm

Energy Dispersive X-ray Spectroscopy (EDAX) was performed to analyze the elemental composition of the wild-type and *tau ^KO^/tau ^KO^* mutant tubules. In the wild-type tubules (Figure 3D and F), the elemental profile was predominantly carbon (C, 83.8%), followed by oxygen (O, 14.1%), with trace levels of phosphorus (P, 1.1%) and calcium (Ca, 0.3%). This composition reflects the organic macromolecules and minor mineral content typical of healthy epithelial tissues (Figure S3, Table 1), whereas significant compositional shifts were observed in *tau ^KO^/tau ^KO^* tubules (Figure 3E and G). The carbon content drastically decreased to 19.8%, whereas the oxygen levels increased to 41.5%, indicating oxidative stress or water retention. Calcium levels markedly increased to 32.3%, potentially due to pathological calcification, while phosphorus levels increased to 3.0%, suggesting altered membrane dynamics or cellular remodeling (Figure S3, Table 1).

This reduction in carbon suggests compromised organic content and cellular degradation, consistent with cytoskeletal abnormalities and impaired function in *tau ^KO^/tau ^KO^* tubules. Elevated oxygen levels could indicate increased oxidative stress or water retention, whereas the drastic rise in calcium suggests disrupted homeostasis and pathological calcification. This increase in phosphorus highlights potential alterations in membrane phospholipids and other phosphorus-containing biomolecules. These findings provide critical insights into the biochemical and structural disruptions occurring in the absence of Tau, further reinforcing its role in maintaining the morphological and functional integrity of MTs.

### 2.4 Early Onset of Tau-Induced Tubule Defects

To determine when Tau depletion begins to impact the morphogenesis of MTs, we examined tubule development during various embryonic stages in *Tau RNAi* and *tau ^KO^/tau ^KO^* embryos. Using immunostaining with the Cut antibody, we visualized the stages of MT development, from their initial specification through proximo-distal (P-D) elongation and circumferential narrowing. In control embryos (*GAL4c42*/Oregon R^+^), MTs followed a coordinated sequence of developmental changes. By embryonic stage 14 (Figure S4C), the tubules underwent significant elongation through convergent extension and showed circumferential narrowing, forming four slender tubular structures with an organized epithelial layer. By stage 16 (Figure S4D), the tubules displayed a mature morphology, including complete elongation, a clear lumen, and well-aligned epithelial cells, consistent with functional differentiation.

In contrast, Tau-deficient embryos exhibited abnormalities starting at stage 14. Both *tau ^KO^/tau ^KO^* (Figure 4B) *and GAL4c42 > UAS tau RNAi* (Figure 4C) embryos showed incomplete elongation and aberrant morphologies. Instead of forming slender, elongated tubules, these embryos displayed distorted and budding structures. The epithelial cells were disorganized, and tubules contained an increased number of cells compared to controls, indicating potential issues with cell rearrangement or regulation of proliferation. By stage 16, these defects became even more pronounced. Tau-deficient tubules (Figure 4E and Figure 4F) remained malformed, with severe elongation and narrowing defects. The epithelial cells lacked proper alignment, and the lumen appeared irregular, failing to achieve the organized structure characteristic of controls (Figure 4D).

**Figure 4.**
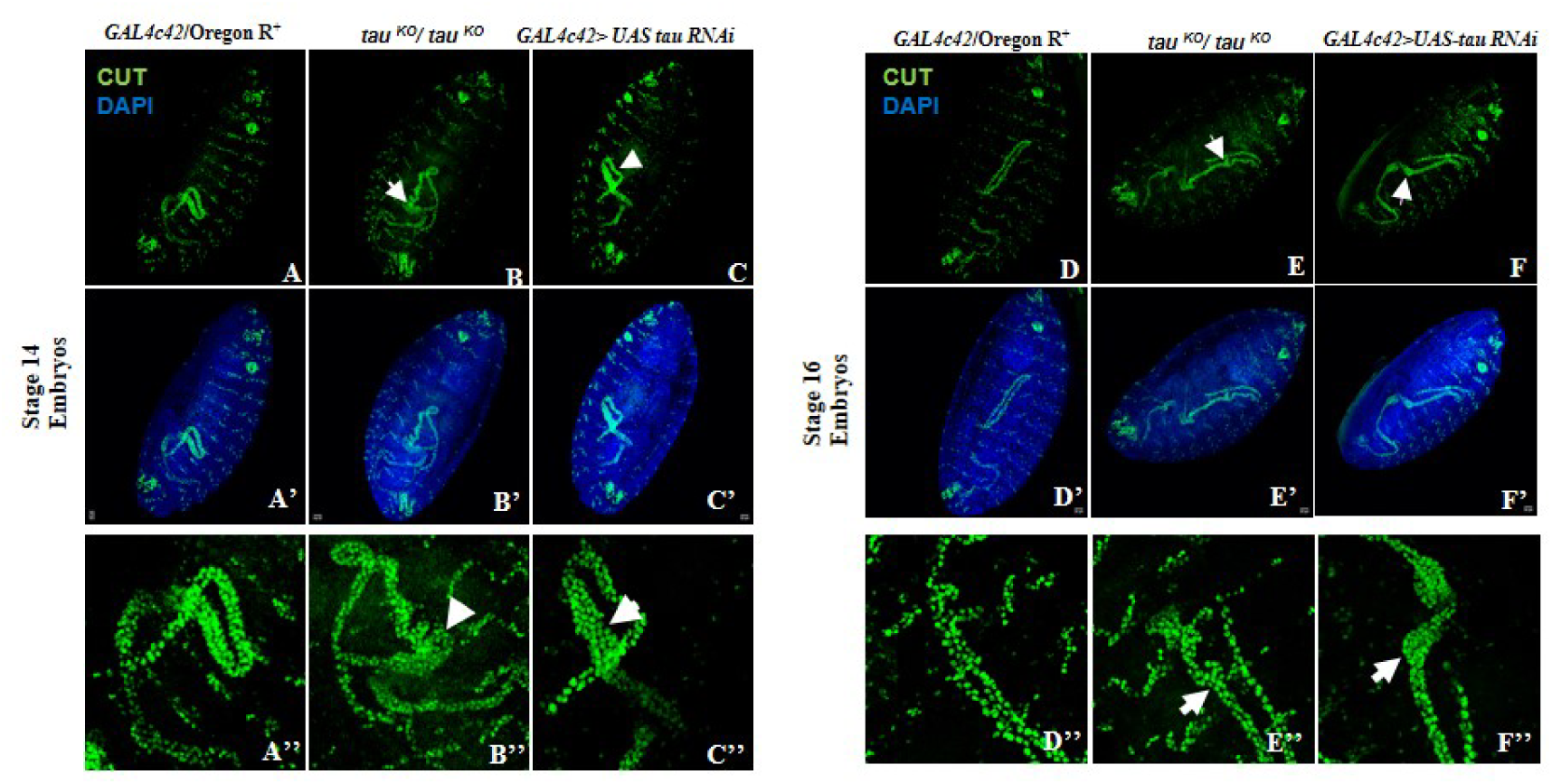
Malpighian Tubule Development is disrupted in *tau ^KO^* and *tau RNAi* conditions from Stage 14 Onwards. In *tau ^KO^* (B, B’ and B”) and RNAi (C, C’ and C”) conditions, initial defects become apparent (white arrow), including disorganized epithelial cells, and irregular lumen formation. Compared to (*GAL4c42*/Oregon R^+^) controls (A), cell alignment is disrupted. A”, B”, C” are magnified images of A, B and C and A’, B’ and C’ shows DAPI staining of the same embryos in A, B, and C. The defects become more pronounced, in stage 16 with severely malformed tubules. The lumen structure is irregular, epithelial organization is compromised (white arrow), and overall morphogenesis is significantly impaired in both *tau ^KO^* (E, E’ and E”) and RNAi (F, F’ and F”) conditions. D’, E’ and F’ shows DAPI staining of the same embryos in D, E and F and D”, E” and F” are magnified images of D, E and F. Scale bar: 20 µm.

These observations highlight Tau’s essential role in driving the key morphogenetic processes that underpin normal MT development, particularly from embryonic stage 14 onward.

### 2.5 Impact of Tau Loss on Tubule Structure: Length, Cell Arrangement, and Cell Count

By the end of embryogenesis, MTs are formed with specific and invariant numbers of PCs and SCs. These cells were arranged in characteristic pattern, with two cells encircling the lumen, a structure achieved through convergent extension movements, and is crucial for tubule function and remains unchanged throughout the lifespan of the fly. In *tau ^KO^/tau ^KO^*mutant conditions, the tubules displayed abnormal morphology, prompting us to measure their length. Comparisons of the anterior and posterior tubules (Figure 5A) revealed a significant increase in the length of *tau ^KO^/tau ^KO^*tubules compared with wild-type controls (Figure 5A).

**Figure 5.**
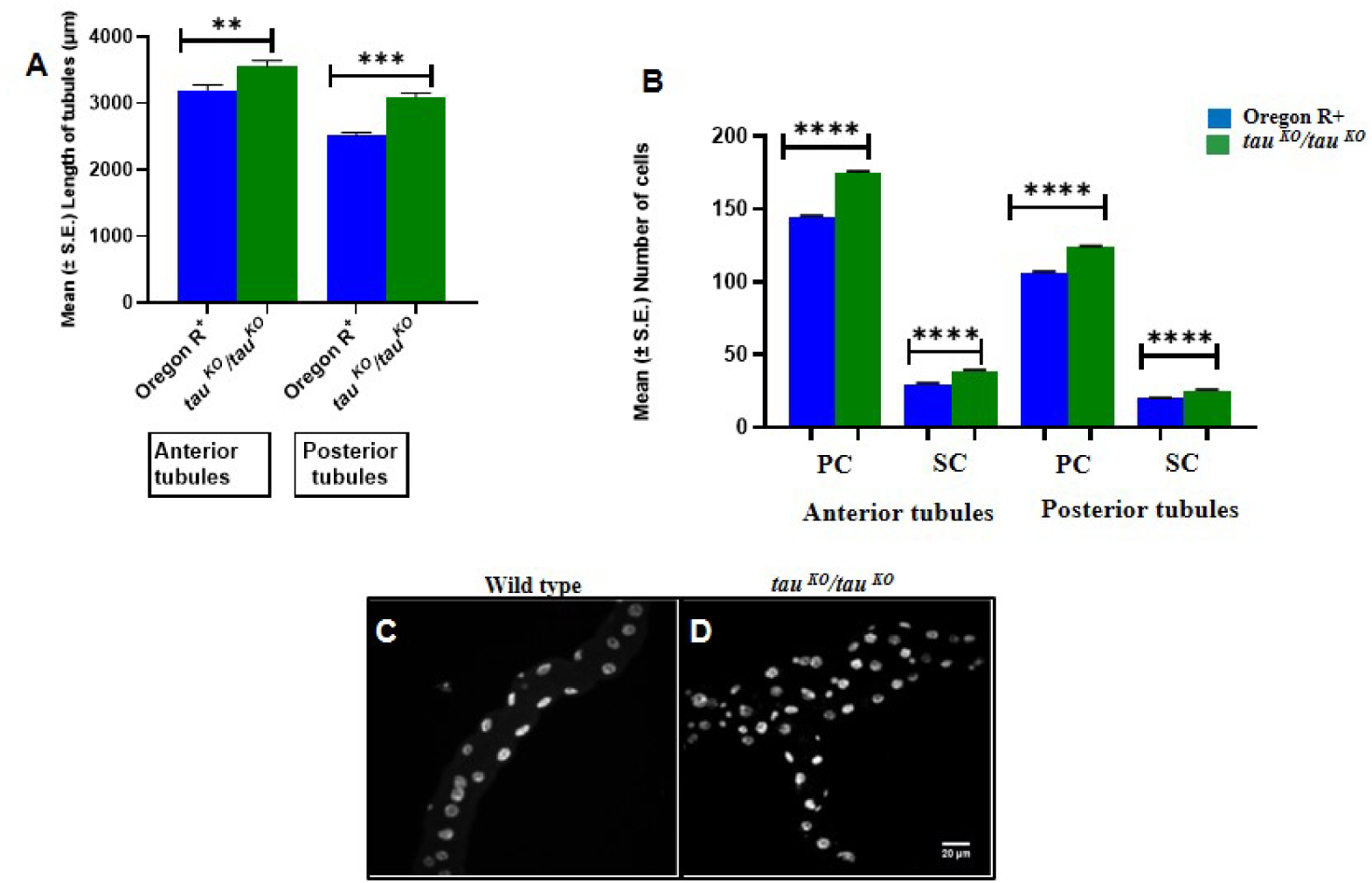
Tau inhibition affects tubule length and SCs number in MTs along with the loss of cellular arrangement. (A) Bar diagram showing significant differences in the mean length (SE±) of anterior and posterior tubules between wild type and mutants (n = 25 pairs of MTs of each genotype). (B) Bar diagram showing mean (SE±) number of PCs and SCs in *tau ^KO^* compared to wild type (n = 25 pairs of MTs of each genotype). The PCs number is more affected compared to SCs in *tau ^KO^*. (C) Wild-type tubules have typical cell arrangement with just two cells around the lumen. This arrangement is lost in *tau ^KO^* (C) and numerous cells present around the lumen. Nuclei are stained by DAPI (Pseudocolor), Images C and D are projections of optical sections obtained by confocal microscope. Statistical analysis was done using t test, asterisk indicates P ≤ 0.05. Scale bar: 20 µm

To determine whether the altered morphology also affected the cellular organization, we examined the arrangement of PCs and SCs. In the wild-type tubules, cells were intercalated at regular intervals, forming a uniform pattern (Figure 5C). However, this pattern was disrupted in *tau ^KO^/tau ^KO^* tubules. Instead of the expected two-cell arrangement encircling the lumen, multiple cells were irregularly clustered together, reminiscent of the morphology observed before convergent extension (Figure 5D). Further analysis of cell numbers showed additional defects in *tau ^KO^/tau ^KO^* tubules. While the total cell count was significantly increased, the most striking observation was an increase in the SC numbers. Wild-type tubules contained approximately 33 SCs in the anterior region and 22 SCs in the posterior region, whereas *tau ^KO^/tau ^KO^* tubules had 45 SCs in the anterior region and 30 in the posterior region (Figure 5B).

These findings highlight the critical role of Tau in maintaining the structural integrity of MTs. It is essential not only to achieve proper tubule length but also to ensure the correct arrangement and balance of cell types during development. The disruption of these processes in Tau-depleted conditions highlights their importance in tubule morphogenesis and function.

### 2.6 Tau-Dependent Cytoskeletal Disorganization and Polarity Defects in Malpighian Tubules

Correct organization of the cytoskeleton is critical for proximo-distal (P-D) elongation and maintenance of MTs morphology. To investigate the impact of Tau loss on cytoskeletal organization, we analyzed the distribution of F-actin and tubulin in third-instar larval tubules. F-actin was visualized by Phalloidin staining. In *GAL4c42*/Oregon R^+^ tubules, F-actin localized primarily to the cell cortex and was enriched at the apical (luminal) surface of tubule cells (Figure 6A, D). In contrast, *tau ^KO^/tau ^KO^* tubules and *GAL4c42> UAS tau RNAi* exhibited a substantial increase in F-actin expression, accompanied by a complete disruption of its organization (Figure 6B, C). The actin filaments appeared dense and diffused throughout the tubule cells with pronounced disarrangement. Notably, actin bundles surrounding stellate cells (SCs) were more compact and exaggerated compared to those in the wild type (Figure 6B, E).

**Figure 6.**
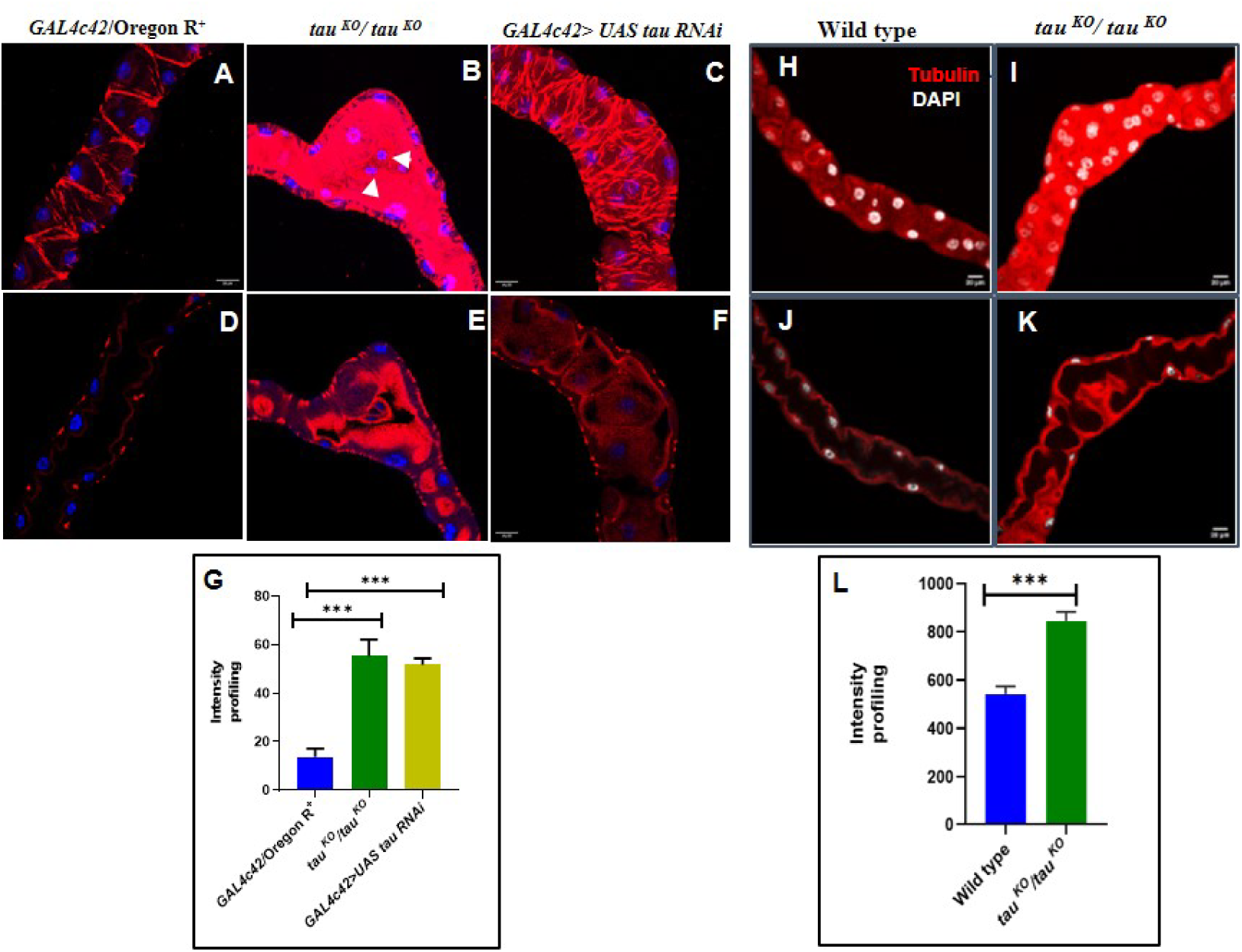
Disruption of cytoskeletal elements in Tau mutants. F-actin organization on cell membrane, in cytoplasm, and at the lumen of MTs of wild-type third instar larvae (A and D) is seen to be disrupted in *tau ^KO^* (B and E) and *GAL4c42>UAS tau RNAi* (C and F). D, E and F are section images of A, B and C, respectively, showing lumen arrangement in tubules. Images A-C are projections of optical sections obtained by confocal microscope. Nuclei are stained by DAPI, scale bar 20 μm. Arrows show enhanced actin accumulation in SCs of *tau ^KO.^* Wild-type tubules showed apical β-tubulin expression (H and J), which is seen to be disrupted in *tau ^KO^* (I and K). Images H and I are projections of optical sections obtained by confocal microscope, images J and K are single section of H and I, respectively. Nuclei are stained by DAPI (Pseudocolor), scale bar 20 μm. (G and L) is a bar diagram showing mean (SE±) intensity profiling of Phalloidin and β-tubulin of Malpighian tubules in *tau ^KO^* and RNAi compared to *GAL4c42*/Oregon R**^+.^** Statistical analysis was done using t test, asterisk indicates P ≤ 0.05.

Tubulin expression, assessed using anti-α-tubulin antibody staining, was also significantly elevated in Tau-depleted tubules compared with *GAL4c42*/Oregon R^+^. In *GAL4c42*/Oregon R^+^ tubules, tubulin showed clear and distinct localization to the luminal region (Figure 6H, J). However, under *tau ^KO^/tau ^KO^* conditions, the tubulin network appeared diffused, with increased expression throughout the cells and a pronounced shift in localization toward the basal membrane (Figure 6 I, K)]. These observations indicate that Tau is required for the proper spatial organization of actin and tubulin in Malpighian tubules during development. To evaluate the apicobasal polarity of MTs cells, we analyzed the localization of Disc large (Dlg), a basolateral membrane protein that is crucial for maintaining epithelial polarity. In *GAL4c42*/Oregon R^+^ tubules, Dlg was distinctly localized to the cell membrane, forming a sharp and uniform pattern (Figure 7A, D). However, under *tau ^KO^/tau ^KO^* and *GAL4c42> UAS tau RNAi* conditions, this precise localization was significantly disrupted. Dlg appeared diffuse and thickened, unlike *GAL4c42*/Oregon R^+^, where it was strictly on the membrane (Figure 7B, C). These observations suggest that the loss of Tau leads to disrupted apicobasal polarity in tubule cells. The compromised polarity likely underlies several structural abnormalities, including weakened cell-cell contacts, the formation of multi-layered cell arrangements, gaps between cells, and the appearance of rounded, protruding cells.

**Figure 7.**
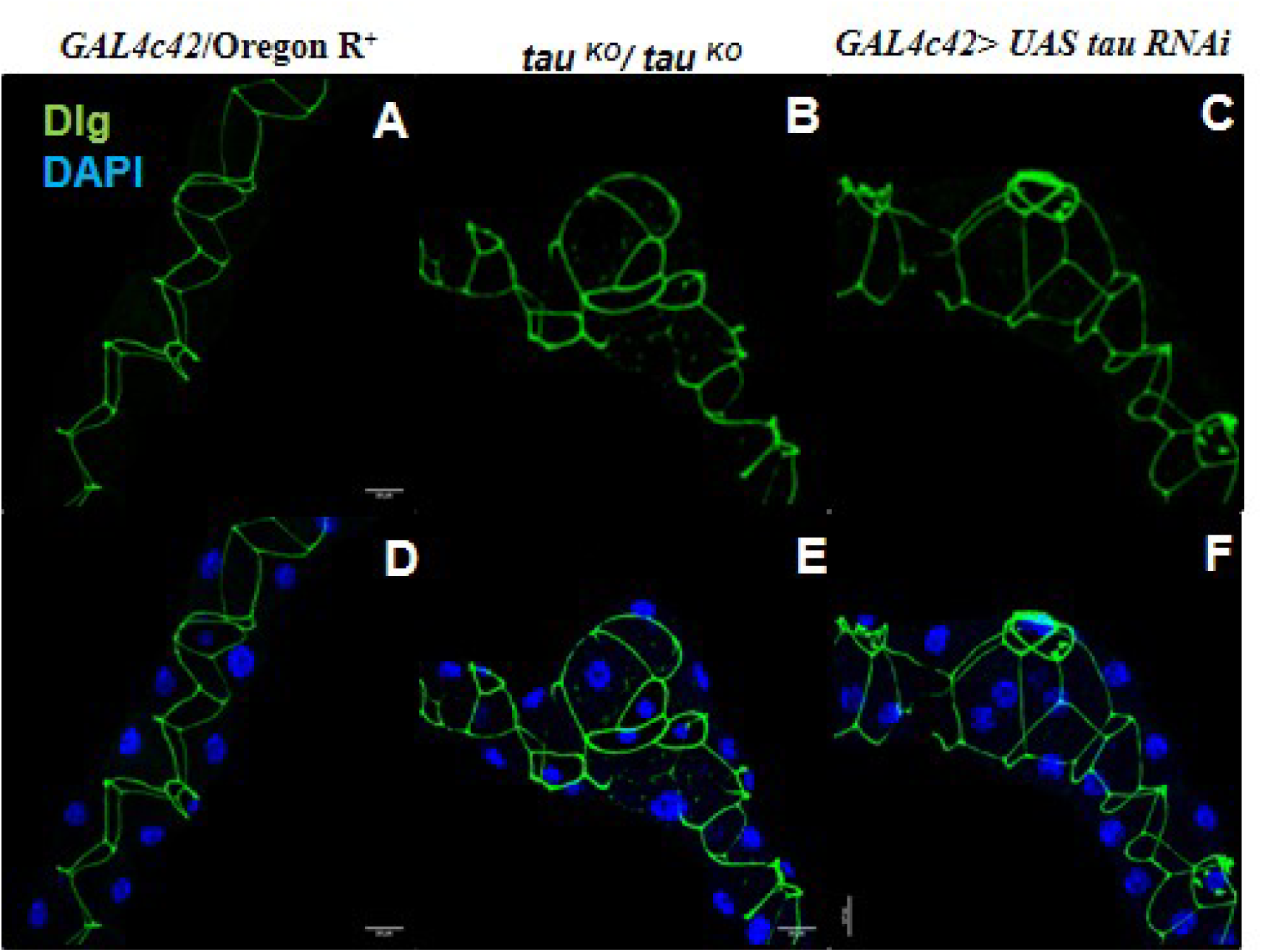
Loss of Tau results in disruption in Disclarge (Dlg) localization. Dlg is localized to the baso-lateral lining of MTs in *GAL4c42*/Oregon R^+^ third instar larvae (A and D), in *tau ^KO^* (B and E) and *tau RNAi* (C and F) Dlg localization is disrupted. Nuclei are stained by DAPI (blue) in D, E, and F. All images are projections of optical sections obtained by confocal microscope, scale bar, 20 μm.

### 2.7 Genetic Crosstalk Between Tau and Rho1 GTPase in Malpighian Tubule Morphogenesis

The intricate process of Malpighian tubule (MT) morphogenesis relies on precise cytoskeletal dynamics and cellular signaling pathways. Our study revealed a critical genetic interaction between Tau, a microtubule-associated protein, and Rho1 GTPase in shaping the morphology and function of tubules. Tau is well known for its role in stabilizing microtubules, while Rho1 GTPase orchestrates actin cytoskeletal remodeling and cellular organization. Under *tau ^KO^/tau ^KO^* conditions, the interplay between these pathways is profoundly disrupted, leading to striking morphological defects and compromised renal function.

Our findings show that loss of Tau results in considerably higher expression of Rho1 (Figure 8B, C) compared to the wild type (Figure 8A), and its downstream effector Rho-associated kinase (Rok), which are central players in actin-microtubule cross-talk. Transverse section images showed that Rho1 localization was more intense on the membrane (Figure 8B’, C’), unlike in the wild type (Figure 8A’). To confirm the role of Rho1GTPase and its association with Tau in MTs tubulogenesis, we downregulated Rho1 expression in *tau ^KO^/tau ^KO^*conditions using *Rho1-72R* (loss of function allele of Rho1) and examined the consequences. In the Rho1-72R background, MTs of *tau ^KO^/tau ^KO^* (n = 40) showed a significant reduction in the severe phenotype compared to that observed in Tau mutants alone. The presence of cysts and buds, which were very prominent in Tau mutants, disappeared in *Rho1-72R/CyO*; *tau ^KO^/tau ^KO^* progenies (Figure 8H), and they closely resembled wild-type MTs (Figure 8F). This observation clearly indicated that the rescue could be due to proper convergent extension resulting from proper cytoskeletal arrangement. To validate this observation, we examined F-actin arrangement and observed that there was a remarkable reduction in the expression level of actin as well as regaining of its arrangement (Figure 7 K) almost similar to that of the wild type (Figure 8I). These results prove that the presence of *Rho1-72R/CyO* in *tau ^KO^/tau ^KO^* rescued their cystic MTs phenotype, strongly suggesting a functional interaction between Rho1GTPase and Tau in MTs.

**Figure 8.**
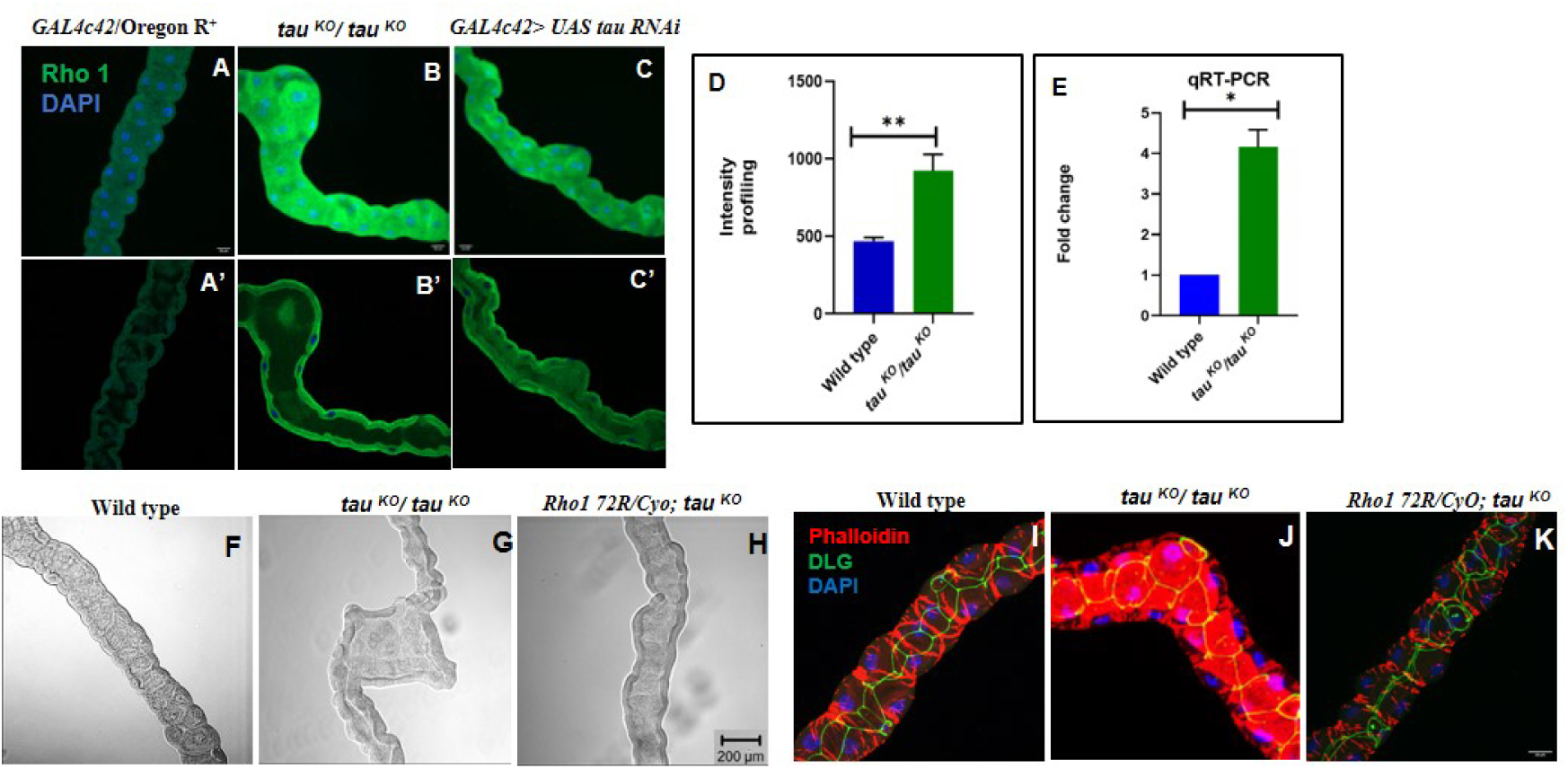
Functional interaction between Tau and Rho1 in Drosophila Malpighian tubules. *GAL4c42*/Oregon R^+^ Malpighian tubules show minimal expression and proper localization of Rho1**(A, A’) while** *tau ^KO^/ tau* ^KO^ (B, B’)and *GAL4c42> UAS tau RNAi* (C,C’) tubules exhibit elevated Rho1 expression and altered localization compared to *GAL4c42*/Oregon R^+^ **(A, A’).** The *tau ^KO^* mutant tubules display defective morphology (G), which is rescued in the *Rho1-72R/CyO; tau ^KO^* background (H), restoring a wild-type-like phenotype (F). **F-actin organization** and **cell arrangement** defects seen in *tau ^KO^* tubules (J) are rescued in the *Rho1-72R/CyO; tau ^KO^* background (K). Nuclei are stained with DAPI (blue). (A-C and I-K) represent projections of confocal optical sections, while (A’-C’) show single optical sections. Scale bar: 20 μm.

### 2.8 Renal Dysfunction in Tau Mutants

Since gross morphological defects and disruption in cellular arrangement were observed in *tau ^KO^/tau ^KO^* mutants, we assessed the functional consequences of MTs, since precise cellular organization is essential for optimal organ function. One of the most prominent differences was observed in the lumen of *tau ^KO^/tau ^KO^* tubules, which exhibited multiple abnormalities, including improper formation, complete absence of lumen, or significantly enlarged lumen size (Figure 9B, C), compared to the well-defined and uniform lumen observed in wild-type tubules (Figure 9A). In the wild type, the maximum width of the lumen was in the range of 54 μm, while in *tau ^KO^/tau ^KO^* mutants it was in the range of 155 μm, which, needless to say, was significantly higher, and the diameter of the narrowest region in the wild type tubule was also significantly less in comparison to the narrowest region of *tau ^KO^/tau ^KO^* mutant tubules (Figure 9D). A single tubule showed variable phenotypes; in some regions, the lumen was not formed, whereas in the adjoining region, it was very much enlarged. There was a noticeable reduction in membrane thickness in the mutants. These defects could be consequential to abnormal actin and tubulin organization, resulting in the loss of lumen formation in the mutants.

**Figure 9.**
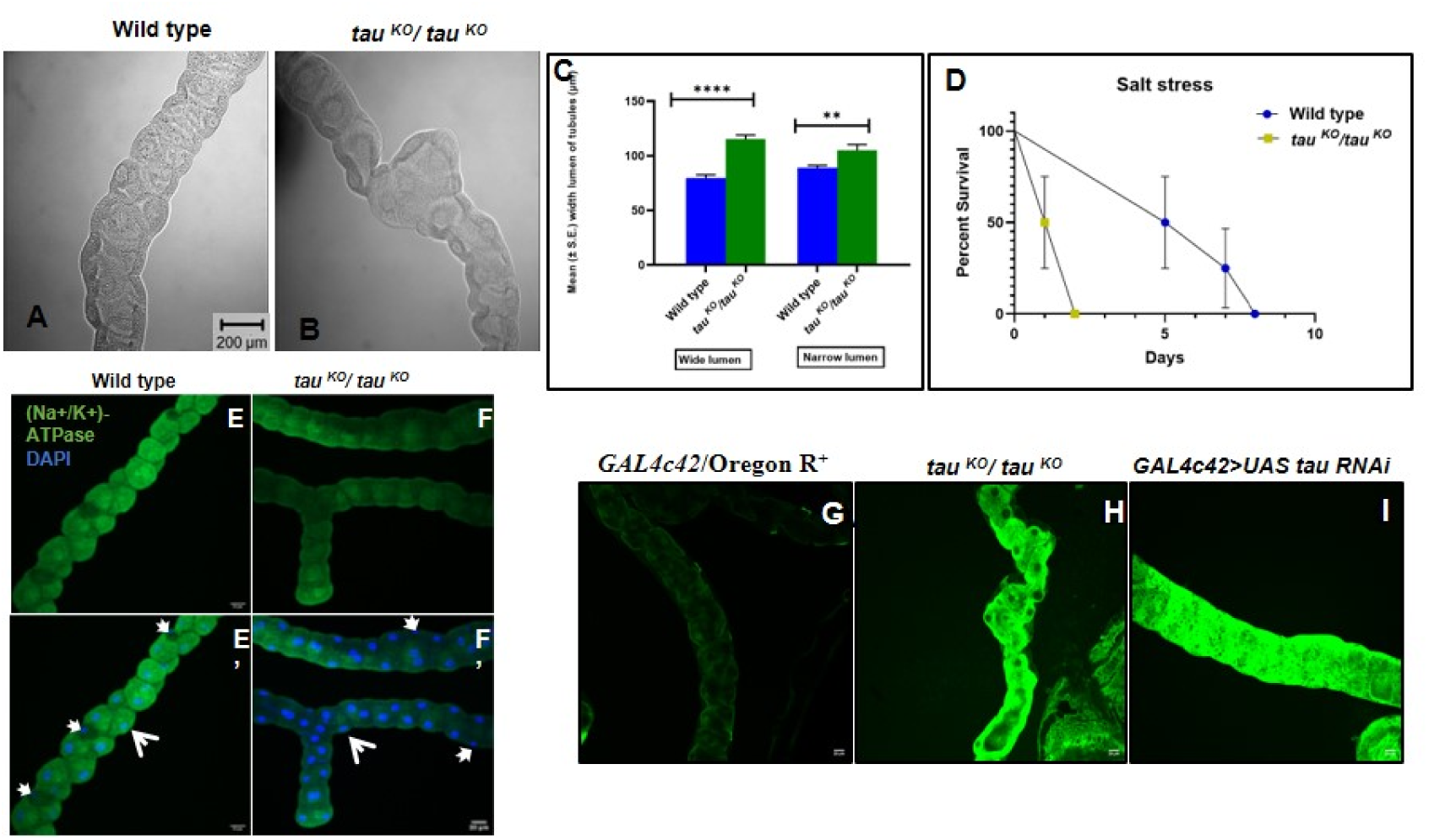
Loss of Tau results in physiological defects in Malpighian tubules. Typical structure of lumen and its size was severely affected by Tau inhibition. DIC images of Malpighian tubules showing lumen architecture in wild type (A) and *tau ^KO^/ tau ^KO^* (B, C). (D) is a bar diagram showing mean (SE±) lumen width of Malpighian tubules in *tau ^KO^* compared to wild type (n = 25 pairs of MTs of each genotype), lumen was categorized into wide and narrow lumen types. Statistical analysis was done using t test, asterisk indicates P ≤ 0.0001 and 0.0085. *tau ^KO^/ tau ^KO^* flies exhibited sensitivity to salt stress (E). Fly populations (wild-type, *tau ^KO^*, n =300, standard deviations are shown) placed in vials containing food with 0.5M NaCl were at a disadvantage and had shorter life span than the wild-type controls. Tau is required for proper localization of ion transporter Na+/K+ ATPase, Tau inhibition causes decreased expression of Na+/K+ ATPase in *tau ^KO^/ tau ^KO^* (F and F’), in comparison to wild type (E and E’). Images E-I are projections of optical sections obtained by confocal microscope. Rhodamine 123 efflux assay shows that the dye is efficiently expelled from the *GAL4c42*/Oregon R^+^ MTs in 3rd instar larval (G) stages. However, the dye accumulated in *tau ^KO^* ^and *GAL4c42>UAS*^ *tau RNAi* 3rd instar larvae (H, I) stage, scale bar, 20 μm.

These morphological defects prompted us to investigate the ability of the tubules to perform essential functions, such as ionic balance and waste transport. Malpighian tubule function was tested by transferring 1^st^ instar larvae to a medium containing 0.5M NaCl. The hypertonic medium acts as a stressor that may be used to reveal diminished Malpighian tubule function. Under high-salt conditions, the survival of the wild-type was greatly impaired, with populations reaching 50% at 5 days and 100% at 8±9 days. Compared to the wild type, *tau ^KO^/tau ^KO^* fly lifespans were shortened, with populations steadily declining from exposure to high salt medium and reaching 50% survival in 1 d and 100% at 2 days, respectively (Fig 9D, n = 300). White concretions, which were present only in the initial segment of the anterior tubules of wild-type larvae (Figure S 5A), were also present in the posterior tubules of *tau ^KO^/tau^KO^* (Figure S 5B). The salt assay, also included an analysis of pupation and eclosion rate to assess developmental outcomes. While a higher proportion of larvae successfully pupated in the wild type, a significant reduction in eclosion was observed in the *tau ^KO^/tau ^KO^* group. This suggests impaired developmental progression in the absence of Tau (Figure S 5C). Under wild-type conditions, the tubules efficiently managed salt excretion, as observed from the normal phenotype in the assay. However, in *tau ^KO^/tau ^KO^* conditions, where salt excretion is impaired, functional deficiency can be directly linked to the downregulation of Na^+^/K^+^ ATPase, as shown by immunostaining.

Na^+^/K^+^ ATPase is a critical enzyme responsible for maintaining the ionic gradient across the epithelial cell membrane, driving secondary active transport processes. Reduced expression of Na^+^/K^+^ ATPase in *tau ^KO^/tau ^KO^* tubules (Figure 9F and F’) disrupts ion transport efficiency, leading to an accumulation of ions within the tubule cells or lumen. This ion imbalance could manifest as defective osmoregulation and a reduced ability to excrete salts efficiently, as demonstrated by the salt assay. Moreover, Na^+^/K^+^ ATPase activity is closely associated with the cytoskeletal integrity. Tau indirectly supports the localization and function of membrane-associated proteins, including ion pumps, by stabilizing microtubules. The cytoskeletal disorganization observed in *tau ^KO^/tau ^KO^*conditions (e.g., increased Rho1 activity and disrupted F-actin organization) might contribute to the mislocalization or instability of Na^+/^K^+^ ATPase in the plasma membrane, further exacerbating ion transport defects.

Similarly, the rhodamine efflux assay highlighted functional impairments in the Tau mutants. *GAL4c42*/Oregon R^+^ tubules efficiently expelled rhodamine within 15 min, indicating robust transporter activity (Figure 9F). However, the *tau ^KO^/tau ^KO^* mutant and *GAL4c42> UAS tau RNAi* tubules exhibited a significant reduction in efflux capacity (Figure 9G, H). This resulted in a pronounced accumulation of rhodamine dye in the 3rd instar larvae, reflecting severe transporter dysfunction. Together, these findings highlight the link between the role of Tau in maintaining structural integrity and its critical function in facilitating key renal processes. The observed morphological and functional defects in the Tau mutant Malpighian tubules reflect a failure to sustain the coordinated cellular activities required for effective renal physiology.

## 3. DISCUSSION

Tau, a microtubule-stabilizing protein, is crucial for maintaining cellular organization and function across diverse epithelial tissues, including the Malpighian tubules (MTs) in *Drosophila*. While extensively studied in the context of neuronal functions (Chan and Bonini, 2000), emerging evidence highlights Tau’s significance in non-neuronal tissues, where it influences epithelial morphogenesis and homeostasis (Lee et.al., 2025). In this study, we demonstrate that Tau is essential for the structural integrity and proper development of MTs, simple epithelial tubules analogous to the mammalian renal system. Loss of Tau leads to disrupted cytoskeletal organization, impaired tubule elongation, altered cell arrangement, and overall morphological defects, highlighting its critical role in convergent extension movements and epithelial tissue integrity (Herczenik and Gebbink, 2008).

Previous studies have established Tau’s involvement in neurodegenerative diseases, where its dysregulation is associated with protein misfolding and aggregation (Hurtle et al., 2024). However, its roles in peripheral tissues remain underexplored. Interestingly, studies in mammalian systems have reported Tau expression in renal podocytes, specialized epithelial cells integral to the kidney filtration barrier (Ávila et al., 2022). Podocytes rely on a highly differentiated architecture with interdigitating processes to maintain filtration efficiency, and the presence of Tau in these cells suggests a conserved role in epithelial function. Although studies in pigs (Garcini et al., 1986) and rats (Gu et al., 1996) have confirmed Tau expression in renal tissues, its precise functional significance in kidney physiology remains largely unknown. Our findings in *Drosophila* MTs provide new insights into Tau’s role in epithelial biology and its broader implications for tissue homeostasis and organogenesis, thereby extending the understanding of protein misfolding pathologies beyond neuronal systems.

Development of MTs begins at embryonic stage 11, with early growth driven by type I cell divisions stimulated by epidermal growth factor (EGF) signaling from the tip cell. This proliferation ensures the formation of a short, fat tube with six to ten cells encircling the lumen (Denholm, 2013; Keller et al., 2000). Morphogenesis continues with cell intercalation, transforming the tubules into their final elongated structure (Skaer, 1989; Broadie et al., 1992). Post-embryonic growth of MTs is achieved through endoreplication in principal cells (PCs) and stellate cells (SCs). Perturbations in key regulatory proteins, such as Tau, have been shown to impair this growth process. Knockout of Tau leads to longer tubules due to increased cell numbers, and defects in cell arrangement. These findings highlight the complex interplay of cytoskeletal remodelling and cell motility, in shaping the morphology and functionality of the MTs.

Actin and tubulin, key cytoskeletal components that interact directly with Tau, play critical roles in maintaining cellular architecture and motility (Garcin and Straube, 2019; Farias et al., 2002). Tau is a well-established microtubule-associated protein that binds to and stabilizes tubulin, promoting microtubule assembly and preventing depolymerization (Barbier et al., 2019). In *tau ^KO^/tau ^KO^* conditions, the loss of this stabilization disrupts the delicate balance between microtubules and the actin cytoskeleton, leading to their mislocalization and disorganization. The observed increase in cortical actin and its disrupted arrangement in Tau-deficient MTs likely impair cell motility and intercalation processes critical for tubule elongation and morphogenesis. Tau’s regulation of actin dynamics may occur indirectly through its interaction with Rho GTPases, such as Rho1, which are upregulated under *tau ^KO^* conditions. Elevated Rho1 activity enhances actin polymerization, contributing to aberrant cortical actin accumulation and impaired cytoskeletal remodelling (Sit and Manser, 2011). Similarly, tubulin mislocalization toward the basal membrane in Tau-deficient MTs highlights the critical role of Tau in microtubule organization. Microtubule instability can disrupt intracellular transport and cellular polarity, resulting in defective epithelial cell alignment and lumen formation (Gudimchuk and McIntosh 2021). Together, these findings highlight that the ability of Tau to coordinate actin and tubulin dynamics is essential for maintaining cytoskeletal integrity, facilitating cell intercalation, and ensuring proper MTs development.

Junctional proteins play a critical role in tissue remodelling and epithelial organization (Blankenship et al., 2006; Bonello et al., 2021), and their mislocalization in *tau^KO^* conditions may interfere with proper cellular patterning and polarity. In Tau-deficient Malpighian tubules (MTs), the mislocalization of junctional proteins such as Discs large (Dlg) disrupts apicobasal polarity, weakening cell-cell adhesion and epithelial integrity. These defects contribute to the disorganized arrangement of PCs and SCs, which are essential for proper tubule morphogenesis. Stabilization of membrane proteins is tightly regulated by their linkage to cytoplasmic scaffolding proteins and the actin/spectrin cytoskeleton (Machnicka et al. 2014). The role of Tau in coordinating actin and tubulin dynamics may indirectly maintain this network, ensuring proper junctional protein localization. The absence of Tau destabilizes these interactions, leading to the redistribution of actin filaments and the misplacement of membrane proteins. In *tau^KO^* tubules, disrupted trafficking and mislocalization of key proteins likely impair epithelial organization and lumen formation (Chaudhary et al., 2018). These defects highlight the critical function of Tau in maintaining cytoskeletal and junctional integrity, which are essential for MT development and homeostasis.

The development of mature MTs is critically dependent on precise regulation of Rho1 GTPase activity, which plays a central role in orchestrating cytoskeletal dynamics and tissue morphogenesis (Denholm et al., 2005). Loss of Tau led to upregulation of Rho1 and its downstream effector, Rho-associated kinase (Rok), both of which are key regulators of actin dynamics. Elevated Rho1 levels were correlated with the observed increase in F-actin expression and disorganization in *tau ^KO^* tubules. Importantly, downregulation of Rho1 in Tau-deficient conditions rescued morphological defects, restoring tubule architecture and cytoskeletal organization. This genetic interaction highlights a critical role for Tau in modulating Rho1-mediated actin-microtubule cross-talk, which is essential for convergent extension movements and proper tubulogenesis.

The functional efficiency of an organ is intricately related to its structural development. Similar to vertebrate nephrons, MTs are tubular organs in which proper cell arrangement and lumen architecture are essential for functionality. Disruptions in these features can result in ion and water imbalances in the hemolymph, leading to lethality or phenotypes resembling polycystic kidney disease (PKD) in humans, characterized by distended nephrons (Piersol GM, 1972). Renal dysfunction was evident from the presence of white deposits in the lumen of Tau mutant tubules, particularly in the posterior pair, suggesting defects in fluid secretion and ion homeostasis. Functional assays further validated this observation. Salt assays demonstrated reduced ion regulation in Tau mutants, which could be attributed to impaired excretory capacity. Additionally, immunostaining for Na^+^/K^+^ ATPase, a key ion transport protein, revealed its mislocalization and reduced expression in Tau*-*deficient tubules, further highlighting disruptions in ion transport mechanisms. Rhodamine staining, used as a tracer for fluid transport, showed significantly reduced dye movement in Tau mutant tubules compared to controls, providing direct evidence for compromised fluid secretion. Collectively, these results suggest that the inability of Tau mutants to maintain proper excretory function arises from disrupted ion transport and fluid regulation, likely due to defects in epithelial polarity and cytoskeletal organization.

Our findings establish Tau as an essential regulator of *Drosophila* MTs development, acting through cytoskeletal stabilization, polarity maintenance, and genetic interactions with Rho1 GTPase. The observed defects in morphology, epithelial organization, and cytoskeletal dynamics highlight the multifaceted role of Tau in tubulogenesis and tissue integrity. This study provides a foundation for exploring the broader roles of Tau in tubular biology and its potential implications in human renal and epithelial diseases.

## 4. EXPERIMENTAL PROTOCOL

### 4.1 Fly culture and stocks

*D. melanogaster* specimens were reared on standardized laboratory medium consisting of 10% yeast, 2% agar, 10% sucrose, 10% autolyzed yeast, 3% nipagin, and 0.3% propionic acid. The cultures were maintained at an ambient temperature of 25 ± 1°C with relative humidity ranging from 50% to 70%, under alternating 12-hour light and dark periods. This study employed the following mutant alleles: *tau ^KO^* mutant flies (Burnouf et al., 2016) kindly supplied by Dr. L. Partridge (Max Planck Institute for Biology of Aging, Cologne, Germany). Dr. J.A.T. Dow (Institute of Biomedical Sciences, University of Glasgow, UK) provided the principal cell-specific Gal4 (c42) and stellate cell-specific Gal4 (c724) drivers. For targeted gene knockdown induction, *UAS-tau RNAi* lines were procured from Bloomington Drosophila Stock Center (#40875). Genetic crosses were established to produce the following experimental genotypes: *GAL4c42 > UAS-tau RNAi*, and *GAL4c724 > UAS-tau RNAi*. Offspring resulting from these crosses were used in the experimental procedures.

### 4.2 Embryo Collection and Preparation

Adult female flies were allowed to oviposit on food plates for 9–24 hours at 24 ± 1°C. Embryos were collected in microcentrifuge tubes, washed with distilled water, and dechorionated by immersion in a 50:50 solution of bleach. Subsequently, embryos were transferred to a 1:1 ml ratio of n-heptane and 4% formaldehyde in 1X Phosphate buffer saline (PBS), and the mixture was agitated gently for 20 min to facilitate fixation. The lower aqueous phase was carefully removed and 2 ml of fresh 100% methanol was added, followed by gentle agitation for 20 s. The supernatant liquids comprising non-devitellinized embryos and empty vitelline sacs were removed, and the embryos that settled at the bottom were retained for further immunohistochemical processing.

## 3. Antibodies and immunohistochemistry

MTs were extracted from healthy third-instar larvae (118–120 h after hatching) in 1X PBS. Tissue samples were fixed in 4% paraformaldehyde for 20 min at ambient temperature, followed by rinsing with 0.1% PBST (1X PBS with 0.1% Triton X-100). A blocking solution containing 0.1% Triton X-100, 0.1% BSA, 10% FCS, 0.1% deoxycholate, and 0.02% thiomersal was applied to the samples for 2 h at room temperature. The sample tissues were kept in the primary antibodies overnight at 4°C. After this incubation period, the samples were washed three times with 0.1% PBST for 20 min each. An additional 2-hour blocking step was performed before incubation with the secondary antibodies. DAPI (1 mg/ml, Molecular Probe) was used to counterstain the tissues for 15 min at room temperature, followed by further washing with 0.1% PBST. For imaging purposes, the samples were mounted in DABCO antifadant (Sigma). The following staining and antibodies were used:

**Stains** : Rhodamine-123 (1 mg/ml), DAPI (1 mg/ml) and Phalloidin 550 (1 :800)

**Primary Antibodies**: Anti-Ct (DSHB, 1:10), Anti-Dlg (DSHB, 1:20), Anti-Rho1 (DSHB, 1:20), Na+/K+ ATPase (DSHB, 1:20), Anti-β tubulin (DSHB, 1:200), Anti-tau (DSHB, 1:20).

## 4. Reverse transcription (RT-PCR)

For RT-PCR analysis, MTs from 50 healthy wandering third instar larvae of different genotypes were dissected. RNA was isolated using the TRIzol method, following the manufacturer’s protocol (Sigma-Aldrich, India) and transcribed into complementary DNA (cDNA). One microgram of cDNA was used as the template and qRT-PCR was performed on a real-time PCR machine (ABI 7500) using appropriate primers and SYBR Green Master Mix with the following primers:

Tau forward: 5′-ACAAAGTTGCAGTGGAACGC-3′ and reverse: 5′-TGGATCTTGATGTCTCCGCC -3′

Rho 1 forward: 5′-CGACGATTCGCAAGAAATTG-3′ and reverse: 5′-CTCGATGTCGGCCACATAAT-3′

The expression data were normalized against the RP49 reference gene and fold changes were calculated relative to the controls.

## 5. Scanning electron microscopy and Energy Dispersive X-ray Spectroscopy (EDAX) spectroscopy

MTs were carefully dissected and fixed in Karnovsky’s fixative for 40 minutes on ice. Following primary fixation, tissues were post-fixed in 1% osmium tetroxide prepared in 0.1 M phosphate buffer (pH 7.4) for 1 h on ice. To ensure complete removal of osmium tetroxide, the samples were washed three times with 0.1 M phosphate buffer (pH 7.4) at 4°C. Subsequently, the tissues were dehydrated in a graded series of cold ethyl alcohol and acetone and critical point dried in liquid carbon dioxide using a Critical Point Dryer (E3000 Series, Quorum Technologies Ltd., UK). Subsequently, the samples were coated with gold using a Sputter Coater (150R ES Plus; Quorum Technologies Ltd., UK). SEM (Zeiss Gemini 3, (560), Germany) and EDAX analyses were used to examine tissue architecture, and the elemental composition and results were recorded using an Intel Pentium IV D computer (Model dx2280 MT, HP Compaq, USA).

### 4.6 Analysis of Malpighian tubules function analysis

#### 4.6.1 Salt stress Assay

Freshly hatched synchronized 1st instar larvae were transferred to either control or salt-enriched food media. Salt stress media were prepared by adding NaCl to standard food media to achieve a final concentration of 0.5 M NaCl. The control medium consisted of standard food without additional salt. Larvae were reared under these conditions at 25 ± 1°C. Larval development was monitored at regular intervals, and survival rates were recorded after exposure to the salt-enriched media. MTs were dissected from 3rd instar larvae (118–120 h post-hatching) exposed to salt stress.

#### 4.6.2 Rhodamine-123 efflux assay

The Malpighian tubules from wandering 3^rd^ instar larvae were dissected in 1X PBS at room temperature and subsequently incubated in Rhodamine-123 solution ((1 µg/ml) for 15 min at 37°C. Immediately after incubation, tissues were chilled for 1-2 mins to stop further dye uptake. Following incubation, the tissues were washed three times with 1X PBS at 37°C, with each wash lasting 10 min, to facilitate the efflux of the dye. After washing, Malpighian tubules were mounted in 1X PBS and scanned for analysis.

## 7. Microscopy and statistical analysis

Confocal imaging was performed using Zeiss LSM 510 and 800 Meta Confocal microscopes using an appropriate laser and filter, and analyzed using ImageJ software. Adobe Photoshop 7.0 was used for intensity quantification and arranging images in the form of panels. All statistical analyses were performed with GraphPad Prism 7.0, using two-tailed unpaired Student’s t-test to assess the significance of the variance in mean values within genotypes. Data are expressed as mean ± standard error of the mean, with three biological replicates (n=3). P values <0.05 (*) were considered statistically significant.

## ACKNOWLEDGEMENTS

We thank Bloomington Drosophila Stock Center, Dr. J. A. T. Dow, Prof. Surajit Sarkar for sharing their fly stocks. We thank Central Discovery Center (CDC) Banaras Hindu University, for the Scanning electron microscope facility. Neha Tiwari was supported by a research fellowship from the University Grant Commission, New Delhi, is highly acknowledged.

## Appendix A. Supplementary material

**Figure S1.**
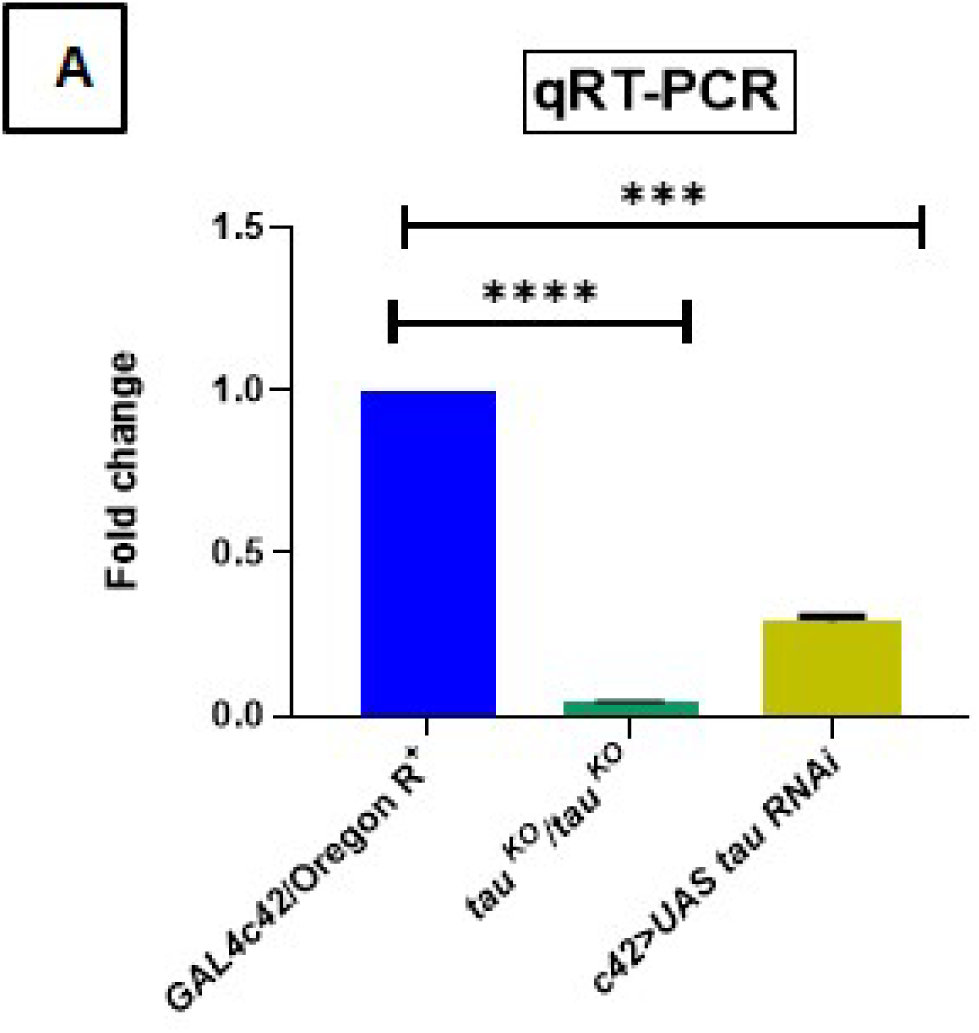
Tau expression **(A)** of *tau ^KO^/tau ^KO^* and *GAL4c42> UAS tau RNAi* compared to the control *GAL4c42*/Oregon R^+^. The bar with * indicates P<0.0001 and P<0.0002.

**Figure S2.**
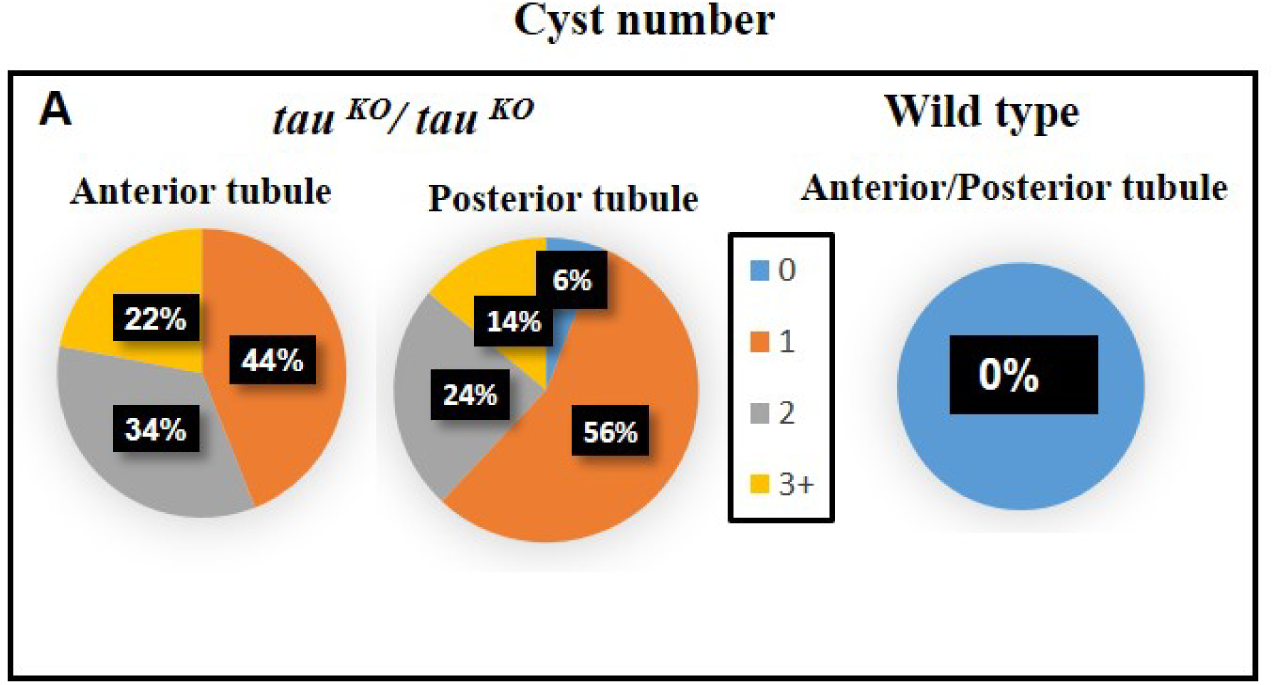
Comparison of cyst numbers in Malpighian tubules under control and *tau ^KO^* conditions. The cysts are categorized based on their count into four groups: 0, 1, 2, and 3+ **(A)**. The percentages of tubules falling into each category are shown for both anterior and posterior tubules.

**Figure S3-Table 1:**
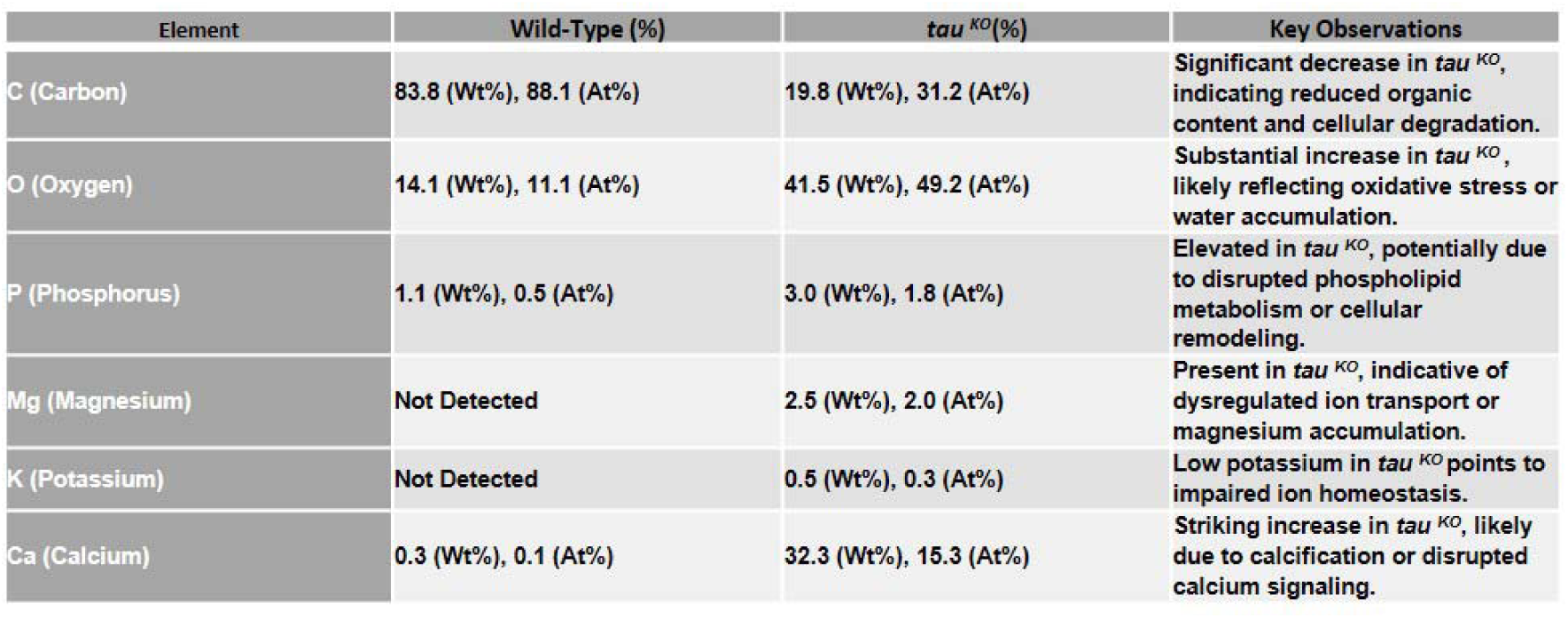
Comparative EDAX analysis of elemental composition in wild-type and *tau ^KO^/tau ^KO^* tubules. The EDAX analysis reveals distinct differences in elemental composition between wild-type and *tau ^KO^/tau ^KO^* Malpighian tubules, shedding light on the physiological and structural changes caused by the loss of Tau.

**Figure S4.**
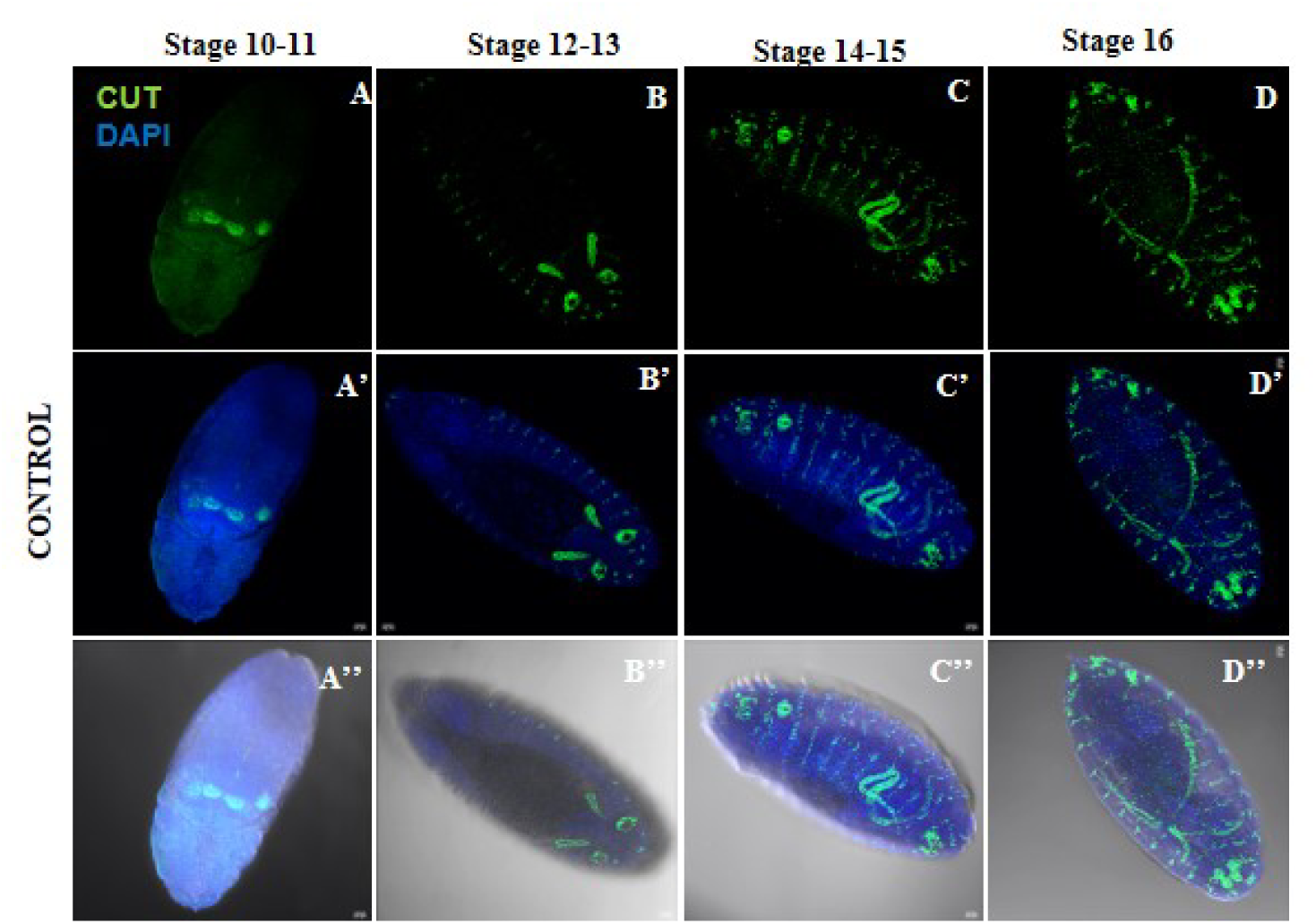
Sequential Development of Malpighian Tubules in Control Embryos: **(A-D)** staining by antibodies against the transcriptional factor Cut of the tubules and central nervous system elements: **(A)** Tubule primordia begin to form as invaginations from the posterior end of the mesoderm. The cells are densely packed, and the initial tube structure is evident; **(B)** Tubules elongate rapidly through convergent extension movements, with clear polarization of epithelial cells. Lumen formation is initiated during this stage. **(C)** Tubules continue elongation, and the lumen becomes more pronounced. Epithelial cells align properly, and the tubules acquire a characteristic shape. **(D)** Tubule morphogenesis is complete, with fully elongated and functional MTs displaying a clear lumen and organized epithelial cell arrangement. A’, B’, C’ and D’ shows DAPI staining of the same embryos in A, B, C and D and A”, B”, C” and D” shows Differential interference contrast images. Scale bar: 20 µm

**Figure S5.**
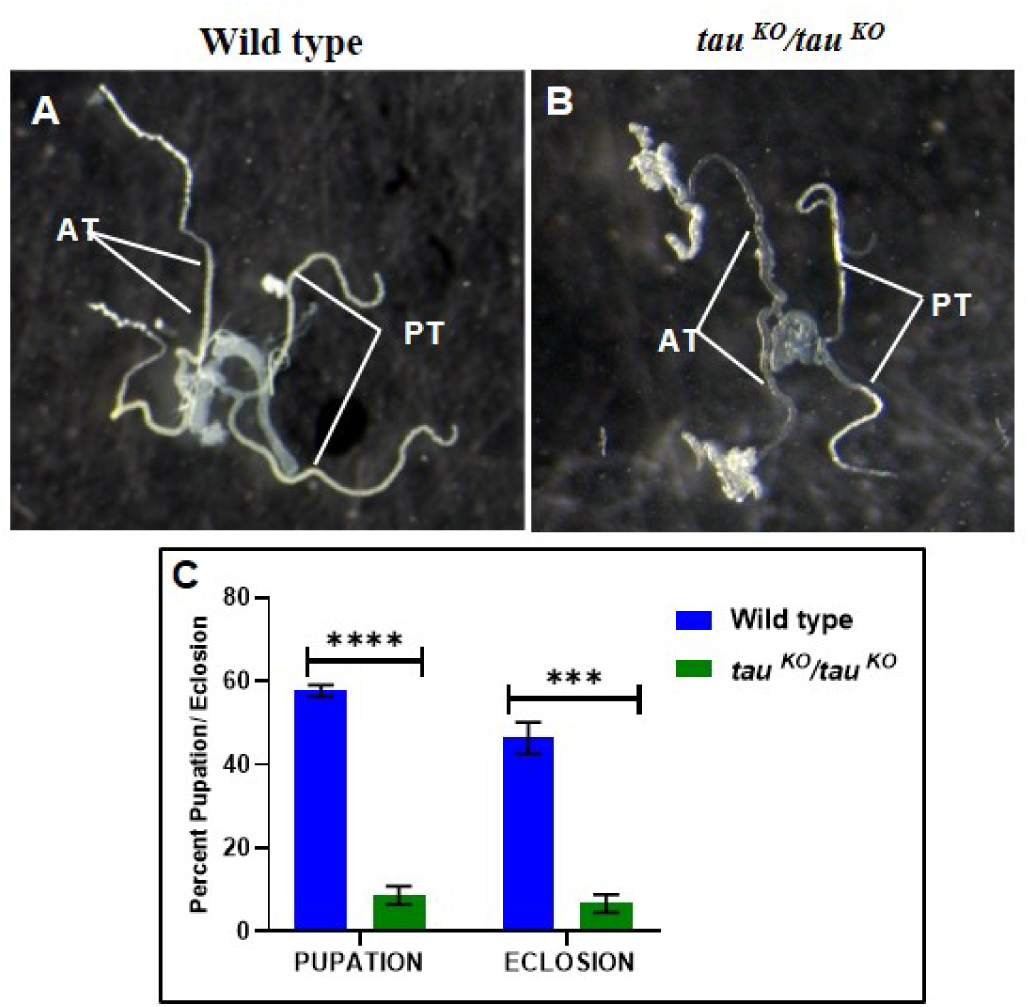
Comparative Analysis of Malpighian Tubule Morphology, Pupation and Eclosion rate in wildtype and *Tau ^KO^* Conditions. **(A-B)** Representative images of Malpighian tubules from wild-type and *tau ^KO^* flies. The posterior tubule of *tau ^KO^* larvae shows abnormal white depositions; a phenotype absent in wild-type tubules. (C) Bar graph showing the number of larvae that pupated and the proportion of pupae that successfully eclosed into adult flies under each condition. Statistical analysis was done using t test, asterisk indicates P ≤ 0.0001 and 0.0009.

## REFERENCES

1. Abrams, EW, Vining, MS and Andrew, DJ (2003). Constructing an organ: The Drosophila salivary gland as a model of tube formation. Trends Cell Biol, 13, 247–254.

2. Arruda, A. P., & Hotamisligil, G. S. (2015). Calcium Homeostasis and Organelle Function in the Pathogenesis of Obesity and Diabetes. Cell metabolism, 22(3), 381–397.

3. Avila J, Lucas JJ, Perez M, Hernandez F (2004). Role of tau protein in both physiological and pathological conditions. Physiol Rev, 84(2), 361–384.

4. Baert L (1978) Hereditary polycystic kidney disease (adult form): a microdissection study of two cases at an early stage of the disease. Kidney international, 13, 519525.

5. Barbier, P., Zejneli, O., Martinho, M., Lasorsa, A., Belle, V., Smet-Nocca, C., Tsvetkov, P. O., Devred, F., & Landrieu, I. (2019). Role of Tau as a Microtubule-Associated Protein: Structural and Functional Aspects. Frontiers in aging neuroscience, 11, 204.

6. Beaven, R., & Denholm, B. (2022). Early patterning followed by tissue growth establishes distal identity in *Drosophila* Malpighian tubules. Frontiers in cell and developmental biology, 10, 947376.

7. Beaven, R. and Denholm, B. (2018). Release and spread of Wingless is required to pattern the proximo-distal axis of Drosophila renal tubules. ELife, 7, e35373.

8. Bertet C, Sulak L, Lecuit T (2004). Myosin-dependent junction remodelling controls planar cell intercalation and axis elongation. Nature, 429(6992), 667–671.

9. Beyenbach, K. W., Skaer, H. and Dow, J. A. (2010). The developmental, molecular, and transport biology of Malpighian tubules. Annu. Rev. Entomol, 55, 351–374.

10. Blankenship JT, Backovic ST, Sanny JS, Weitz O, Zallen JA (2006). Multicellular rosette formation links planar cell polarity to tis sue morphogenesis. Dev Cell, 11(4), 459–470. 39.

11. Bonello T, Aguilar-Aragon M, Tournier A, Thompson BJ (2021). A picket fence function for adherens junctions in epithelial cell polarity. Cells Dev, 203719, 40.

12. Broadie K, Skaer H, Bate M (1992). Whole-embryo culture of *Drosophila*: development of embryonic tissues in vitro. Roux Arch Dev Biol, 201(6), 364–375.

13. Brooke, BS, Karnik, SK and Li, DY (2003). Extracellular matrix in vascular morphogenesis and disease: structure versus signal, Trends Cell Biol, 13, 51–56.

14. Bunt, S., Hooley, C., Hu, N., Scahill, C., Weavers, H. and Skaer, H. (2010). Hemocyte-secreted type IV collagen enhances BMP signaling to guide renal tubule morphogenesis in Drosophila. Dev. Cell, 19, 296–306.

15. Burnouf, S., Grönke, S., Augustin, H., Dols, J., Gorsky, M. K., Werner, J., Kerr, F., Alic, N., Martinez, P., & Partridge, L. (2016). Deletion of endogenous Tau proteins is not detrimental in Drosophila. Scientific reports, 6, 23102.

16. Chan, H., Bonini, N (2000). *Drosophila* models of human neurodegenerative disease. Cell Death Differ, 7, 1075–1080.

17. Chaudhary, A. R., Berger, F., Berger, C. L., & Hendricks, A. G. (2018). Tau directs intracellular trafficking by regulating the forces exerted by kinesin and dynein teams. *Traffic (Copenhagen*, Denmark), 19(2), 111–121.

18. Dell, KM, Watkins, RSL and Avner, ED (2004). Polycystic kidney disease. Pediat Nephrol, 5, 675–699.

19. Denholm B, Brown S, Ray RP, Ruiz-Gomez M, Skaer H, Hombría JC (2005). Crossveinless-c is a RhoGAP required for actin reorganisation during morphogenesis. Development, 132(10), 2389–2400.

20. Denholm B (2013). Shaping up for action: the path to physiological maturation in the renal tubules of Drosophila. Organogenesis, 9(1), 40–54.

21. Denholm, B., Hu, N., Fauquier, T., Caubit, X., Fasano, L. and Skaer, H. (2013). The tiptop/teashirt genes regulate cell differentiation and renal physiology in Drosophila. Development, 140, 1100–1110.

22. Dow, J. A. and Romero, M. F. (2010). Drosophila provides rapid modeling of renal development, function, and disease. *Am. J. Physiol*, Renal. Physiol, 299, F1237–F1244.

23. Dow, J.A.T. (2012). The versatile stellate cell-more than just a space-filler. J. Insect Physiol. 58, 467–472.

24. Dressler, GR. (2002). Tubulogenesis in the developing mammalian kidney. Trends Cell Biol, 12, 390 395.

25. Farias, G. A., Muñoz, J. P., Garrido, J., & Maccioni, R. B. (2002). Tubulin, actin, and tau protein interactions and the study of their macromolecular assemblies. Journal of cellular biochemistry, 85(2), 315–324.

26. Garcin, C., & Straube, A. (2019) Microtubules in cell migration. *Journal of Biochemistry*, EBC20190016.

27. Goedert M, Spillantini MG (2019). Ordered assembly of tau protein and neurodegeneration. Adv Exp Med Biol, 1184, 3–21.

28. Gotz J, Halliday G, Nisbet RM (2019). Molecular pathogenesis of the tauopathies. Annu Rev Pathol,14, 239–261

29. Grantham JJ, Geiser JL, Evan AP (1987). Cyst formation and growth in autosomal dominant polycystic kidney disease. Kidney international, 31, 11451152.

30. Gu Y, Oyama F, Ihara Y (1996) Tau is widely expressed in rat tissues. J Neurochem, 67(3), 1235–1244.

31. Gudimchuk, N.B., McIntosh, J.R. (2021). Regulation of microtubule dynamics, mechanics and function through the growing tip. Nat Rev Mol Cell Biol, 22, 777–795.

32. Hatton-Ellis, E., Ainsworth, C., Sushama, Y., Wan, S., Vijayraghavan, K. and Skaer, H. (2007). Genetic regulation of patterned tubular branching in Drosophila. Proc. Natl. Acad. Sci. USA, 104, 169–174.

33. Herczenik, E., & Gebbink, M. F. (2008). Molecular and cellular aspects of protein misfolding and disease. FASEB journal: official publication of the Federation of American Societies for Experimental Biology, 22(7), 2115–2133.

34. Hurtle, B., Donnelly, C.J., Zhang, X. et al. (2024). Live-cell visualization of tau aggregation in human neurons. Commun Biol, 7, 1143.

35. Jung, A. C., Denholm, B., Skaer, H. and Affolter, M. (2005). Renal tubule development in Drosophila: A closer look at the cellular level. J. Am. Soc. Nephrol, 16, 322–328.

36. Keller R, Davidson L, Edlund A, et al. (2000). Mechanisms of convergence and extension by cell intercalation. Philos Trans R Soc Lond B Biol Sci, 355(1399), 897–922. 33.

37. Kim, Y., Lee, S., and Han, C. (2019). Tau protein: Mechanisms of toxicity and its potential role in renal function. Biology, 8(3), 67.

38. Lee, J., Kim, D., Cha, S. J., Lee, J. W., Lee, E. Y., Kim, H. J., & Kim, K. (2025). Tau reduction impairs nephrocyte function in Drosophila. BMB reports, 6221.

39. Li, Z., Liu, S. and Cai, Y. (2014). Differential Notch activity is required for homeostasis of malpighian tubules in adult Drosophila. J. Genet. Genomics, 41, 649–652.

40. Li, Z., Liu, S. and Cai, Y. (2015). EGFR/MAPK signaling regulates the proliferation of Drosophila renal and nephric stem cells. J. Genet. Genomics, 42, 9–20.

41. Liu, C., Song, X., Nisbet, R., and Götz, J. (2021). Tau and Beyond: The Role of Tau in Organ Function and Dysfunction. Journal of Molecular Biology, 433(6), 166735.

42. Machnicka, B., Czogalla, A., Hryniewicz-Jankowska, A., Bogusławska, D. M., Grochowalska, R., Heger, E., Sikorski, A.F. Spectrins (2013). A structural platform for stabilization and activation of membrane channels, receptors and transporters. Biochimica et Biophysica Acta (BBA), Biomembranes.1838(2), 620–634.

43. Millet-Boureima, C., Porras Marroquin, J. and Gamberi, C. (2018). Modeling renal disease ‘On the Fly’. BioMed Res. Int, 5697436.

44. de Garcini, E. M., Díez, J., & Avila, J. (1986). Quantitation and characterization of tau factor in porcine tissues. Biochimica et Biophysica Acta (BBA)-General Subjects, 881(3), 456–461.

45. Piersol GM (1927). Polycystic disease of the kidney. Trans Am Climatol Clin Assoc, 43, 221–231.

46. Singh, S. R., Liu, W. and Hou, S. X. (2007). The adult Drosophila malpighian tubules are maintained by multipotent stem cells. Cell Stem Cell, 1, 191–203.

47. Sit, S.T., & Manser, Ed (2011). Rho GTPases and their role in organizing the actin cytoskeleton. J Cell Sci, 124 (5), 679–683.

48. Skaer H (1989). Cell division in Malpighian tubule development in *D. melanogaster* is regulated by a single tip cell. Nature, 342, 566–569.

49. Skaer, H. (1996). Cell proliferation and development of the Malpighian tubules in Drosophila melanogaster. Exp. Nephrol, 4, 119–126.

50. Takashima, S., Paul, M., Aghajanian, P., Younossi-Hartenstein, A. and Hartenstein, V. (2013). Migration of Drosophila intestinal stem cells across organ boundaries. Development, 140, 1903–1911.

51. Uv, A, Cantera, R and Samakovlis, C (2003). Drosophila tracheal morphogenesis: intricate cellular solutions to basic plumbing problems, Trends Cell Biol, 13, 301–309.

52. Vallés-Saiz, L., Peinado-Cahuchola, R., Ávila, J. et al. (2022). Microtubule-associated protein tau in murine kidney: role in podocyte architecture. Cell. Mol. Life Sci, 79, 97.

53. Wang, J., Kean, L., Yang, J., Allan, A. K., Davies, S.A., Herzyk, P and Dow, J.A. (2004). Function-informed transcriptome analysis of Drosophila renal tubule. Genome Biol, 5, R69.

